# Mechanistically distinct BER processes are essential for the tolerance of exogenous 5hm-dC and enzymatic oxidative demethylation

**DOI:** 10.64898/2026.08.21.746167

**Authors:** Job Smink, Hannah K. Webb, Eleni Stefoudi, Ross Hill, Michael A. Terzidis, Gerry P. Crossan, Juan I. Garaycoechea

## Abstract

Epigenetic information is transmitted through covalent DNA modifications that regulate chromatin structure and gene expression. 5-Hydroxymethylcytosine (5hmC) is a key epigenetic intermediate generated during TET-mediated oxidative demethylation of 5-methylcytosine. Whether 5hmC itself is intrinsically genotoxic remains unclear.

Here, we combine genome-wide CRISPR loss-of-function screens, isogenic knockouts and mass spectrometry to systematically compare the cellular consequences of exogenous 5-hydroxymethyl-2′-cytidine (5hm-dC) exposure and endogenous oxidative demethylation in mammalian cells. We find that exogenous 5hm-dC causes toxicity, mediated by deamination to 5-hydroxymethyl-2′-deoxyuridine (5hm-dU), followed by excision by the glycosylase SMUG1, which generates base excision repair (BER) intermediates that compromise cell viability. In contrast, toxicity associated with enzymatic oxidative demethylation is primarily driven by TDG-dependent excision of oxidized methylcytosine derivatives. Despite these distinct initiating events, both genotoxins converge on a critical requirement for DNA polymerase β (POLβ), indicating that efficient BER completion is essential to mitigate cytotoxic repair intermediates.

## Introduction

DNA methylation is a key epigenetic mechanism by which cells convey information without altering the genetic code. It plays essential roles in development, genomic imprinting, cell differentiation, chromatin structure, and the regulation of gene expression. DNA methylation occurs through the addition of a methyl group to cytosine (C), generating 5-methylcytosine (5mC), predominantly in a CpG dinucleotide context. This modification is established *de novo* by methyltransferases DNMT3A and DNMT3B and subsequently maintained through cell divisions by DNMT1. As the most abundant DNA modification, 5mC is often referred to as the “fifth base” (1).

Importantly, DNA methylation is highly dynamic; active and passive demethylation restore cytosine residues, and this balance is essential for proper mammalian development. Passive DNA demethylation requires the inhibition of DNMT1-mediated maintenance methylation or DNMT3A/B-mediated *de novo* methylation (2, 3), which is achieved through inhibition of DNMT activity by either physical exclusion of DNMT proteins or downregulation of *Dnmt* expression(4). Active demethylation happens via oxidation of 5mC and subsequent removal by the base excision repair (BER) pathway.

During active demethylation, the ten-eleven translocation (TET) family of dioxygenases— TET1, TET2, and TET3—oxidise 5mC to 5-hydroxymethylcytosine (5hmC) and have the ability to further oxidise to 5-formylcytosine (5fC) and 5-carboxylcytosine (5caC) (5, 6). These modified bases are not substrates for DNMT1 and therefore methylation is lost by loss of maintenance and passive dilution. However, 5fC and 5caC are substrates for BER, specifically for thymine DNA glycosylase (TDG) (and NEIL1 for 5caC (7)), resulting in apurinic/apyrimidinic (AP) sites (8, 9). After removal of the base, AP endonuclease 1 (APE1) makes an incision in the backbone, followed by gap filling by the BER-specific DNA polymerase β (POLβ) in short-patch repair, or by replicative polymerases δ and ε in long-patch repair (10). If not resolved, the generated AP sites may result in genotoxic single-strand breaks (SSBs), double-strand breaks (DSBs) and genome instability. Indeed, TDG-mediated excision of 5fC and 5caC has recently been shown to generate SSBs in neurons, which are resolved by short-patch or long-patch BER depending on cellular context (11, 12), leading to hotspots of DNA synthesis associated with mutations(13). These observations raise the possibility that oxidative DNA demethylation is not merely a passive epigenetic process but may itself represent an intrinsic source of DNA damage.

Recent work has also revealed the potential risks associated with 5hmC, a key intermediate in active DNA demethylation. While 5hmC is generally considered an epigenetic mark, emerging evidence suggests that its metabolic byproducts may compromise genome stability. In particular, exogenously supplied 5-hydroxymethyl-2′-deoxycytidine (5hm-dC) can be deaminated within the nucleotide pool to generate 5-hydroxymethyl-deoxyuridine (5hm-dU) (14, 15). Upon incorporation into DNA, deaminated bases such as 5hmU and uracil may slow the replication fork themselves or be excised, leading to the formation of AP sites that require efficient repair via BER (16, 17). Consistent with this, accumulation of such lesions has been linked to replication fork instability, particularly in homologous recombination and Fanconi anaemia–deficient cells (14, 15, 18). These findings suggest that intermediates of DNA demethylation may be a source of modified nucleosides and contribute to replication stress. Supporting this notion, it has been proposed that 5hm-dC could re-enter the nucleotide pool following oxidative DNA demethylation (15); however, how 5hmC is removed from DNA and whether enzymatic oxidative demethylation is a significant source of 5hm-dU remains unclear. In addition, the potential for direct deamination of 5hmC within DNA is not well understood. Available evidence suggests that cytidine deaminases such as APOBEC and AID exhibit limited activity toward 5hmC in single-stranded DNA (19). Furthermore, other oxidized cytosine derivatives may present additional challenges to genome stability, as 5fC has been shown to form DNA–protein cross-links that can impede DNA replication (20). Together, these observations highlight our incomplete understanding of the fate of epigenetic nucleosides and the mechanisms by which they contribute to genome instability.

Notably, much of our current understanding of 5hmC-associated genotoxicity derives from exogenous systems in which 5hm-dC is directly supplied to cells. While informative, these approaches bypass the endogenous dynamics of oxidative DNA demethylation, leaving a critical gap in our understanding of how this process impacts genome stability under physiological conditions. Here, we systematically define the mechanisms that protect cells from exogenous 5hm-dC and endogenous enzymatic oxidative DNA demethylation, and investigate the crosstalk between these pathways. Using a genome-wide CRISPR/Cas9 screen, we first identify factors that safeguard genome integrity in response to exogenous 5hm-dC. In parallel, we developed an inducible system that drives high levels of oxidative DNA demethylation and directly compare genetic requirements in an isogenic system. Finally, we couple our treatments and genetic models with direct quantification of epigenetic base modifications by LC-MS. Together, our findings establish the routes of epigenetics-induced DNA damage and demonstrate that, although both exogenous and endogenous processes converge on BER dependency, they originate from fundamentally different molecular intermediates.

## Results

### Loss of DNA repair factors hypersensitise 32D cells to exogenous 5hm-dC

5hmC is an important epigenetic DNA modification. However, the cellular factors required for processing and tolerating 5hmC remain poorly defined. Therefore, we sought to identify pathways required for cellular resistance to 5hmC. We performed a genome-wide unbiased CRISPR/Cas9 sensitivity screen using addition of exogenous 5hm-dC in mouse 32D cells (**Fig. 1A**). In short, wildtype Cas9-expressing 32D cells were transduced with a genome-wide sgRNA library and exposed to 5hm-dC. Samples collected before treatment (T0) and after the second and third treatment rounds (T1/T2) were analysed by next-generation sequencing to identify genes required for survival. 32D cells were selected because they are rapidly dividing mouse bone marrow-derived cells with an intact TP53 response, enabling the study of TP53-dependent 5hm-dC sensitivity and translation to *in vivo* mouse models.

**Figure 1.**
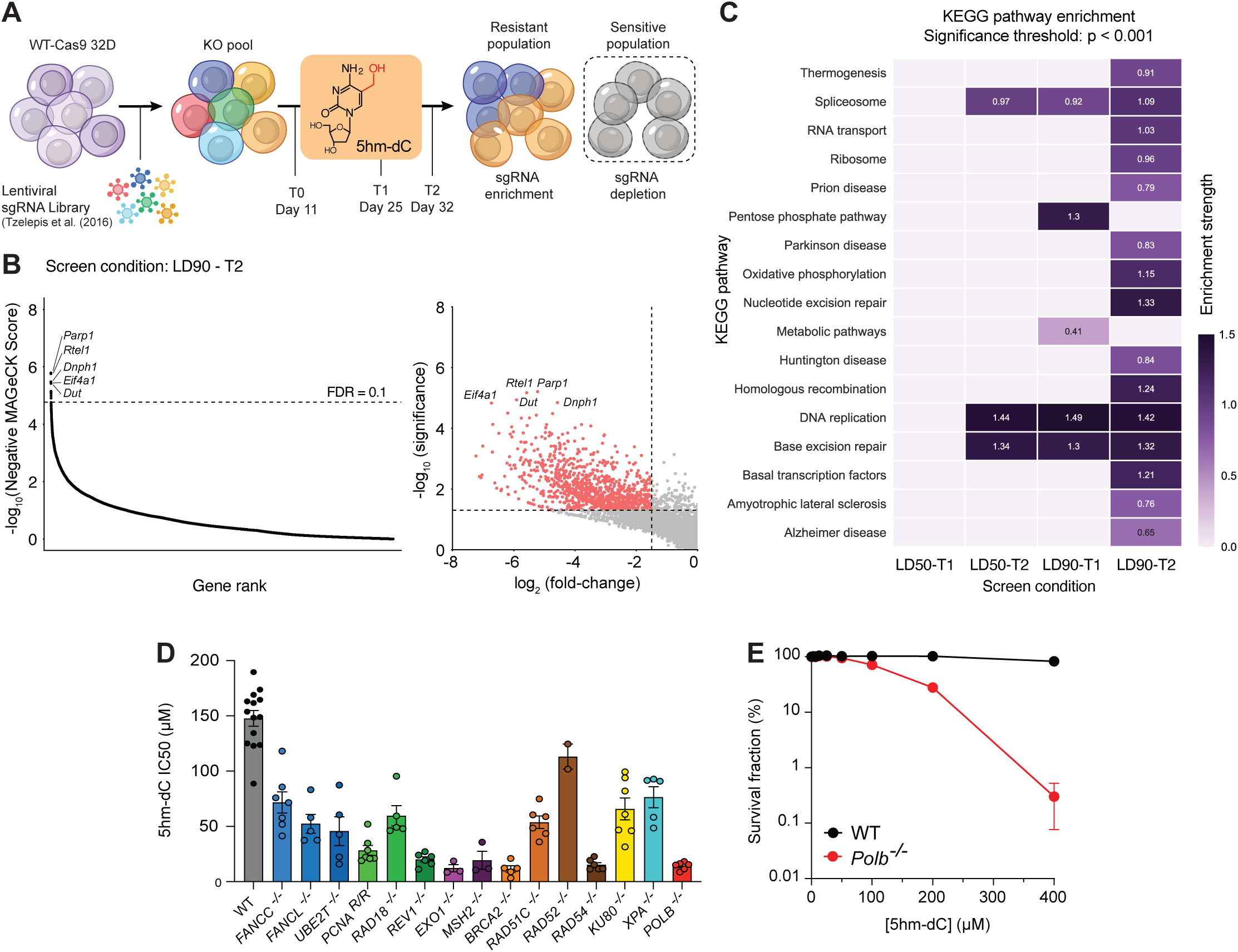
Genome-wide CRISPR/Cas9 screening identifies DNA repair pathways as a key determinant of cellular tolerance to exogenous 5hm-dC. **A)** Schematic overview of the genome-wide CRISPR/Cas9 sensitivity screen in 32D mouse myeloid progenitor cells. Cas9-expressing 32D cells were transduced with a whole-genome sgRNA library and exposed to 5-hydroxymethyl-2′-deoxycytidine (5hm-dC). Samples were collected prior to treatment (T0) and after two and three sequential treatments (T1 and T2). sgRNA representation was determined by next-generation sequencing to identify depleted and enriched guides. **B)** MAGeCK analysis of sgRNA abundance comparing untreated and 5hm-dC-treated cells under LD90 conditions at T2. Genes were ranked based on statistical significance, fold change, MAGeCK score and false discovery rate. The left column shows screen results ranked by MAGeCK score, indicating the FDR cutoff (FDR < 0.1). The right column shows volcano plots with the significance scores from MAGeCK analysis. The top five ranked hits according to MAGeCK score for each condition are labelled. **C)** KEGG pathway enrichment analysis of significantly depleted genes (*P* < 0.001, log2 fold change < −1.5). Multiple DNA repair pathways were enriched, including base excision repair, nucleotide excision repair, homologous recombination **D)** Quantification of 5hm-dC IC50 values in wild-type (WT) and DNA repair–deficient DT40 cells. Data points represent mean ± s.e.m. from n=3-14 independent biological replicates, each comprising two technical replicates. Statistical significance was determined using an unpaired t-test (*P* < 0.001). **E)** Survival curve of WT and *Polb^−/−^* cells following increasing 5hm-dC treatment, measured by fluorescent viability assay. Data points represent mean ± s.e.m. of n=3 biological repeats (n=3 technical repeat each)

Using a model-based analysis of genome-wide CRISPR/Cas9 Knockout (MAGeCK) algorithm and integrating statistical significance (P value), fold-change (population depletion), and false discovery rate (FDR) (see **Methods**), we identified differential sgRNA between untreated and 5hm-dC-treated conditions. (**Fig. 1B**). Ranking genes by essentiality (negative MAGeCK score) across all conditions revealed three recurrent top five hits poly(ADP-ribose) polymerase 1 (*Parp1*), Regulator of telomere length 1 (*Rtel1*), 5-hydroxymethyl-dUMP N-hydrolase (*Dnph1*) and dUTPase (*Dut*) appearing in multiple screens (**Fig. 1B, Supplementary Fig. 1A**). In multiple conditions, these genes showed strong depletion (log2 fold change < −4) with high statistical significance (*P* < 0.0001, FDR < 0.1). *Dnph1* had been identified previously, providing confidence in the validity of our screening results (15). In addition, we utilized the KEGG (Kyoto Encyclopaedia of Genes and Genomics) classification system to explore the broader biological pathways involved in 5hm-dC processing with two different stringency conditions (**Fig. 1C, Supplementary Fig. 1B**). We found there was enrichment of several DNA repair pathways: including BER, nucleotide excision repair (NER), homologous recombination (HR), non-homologous end joining (NHEJ) and Fanconi anaemia (FA) pathway. Both BER and NER were enriched across all conditions, consistent with their central roles in safeguarding the genome against modified bases and diverse base lesions.

To validate the results of the screen, we determined DNA repair requirements for 5hm-dC tolerance were conserved in another vertebrate cell line. We exposed a panel of DNA repair knockout DT40 mutants to increasing concentrations of 5hm-dC and measured cell survival. In agreement with previous analyses, loss of all DNA repair factors increased sensitivity to elevated exogenous 5hm-dC. (**Fig. 1D, Supplementary Fig. 1C**). Specifically, loss of HR and BER factors (BRCA2, RAD54, POLB) increased sensitivity and indicates a conserved mechanism across different vertebrate species. Interestingly, factors in the MMR pathway (*MSH2*, *EXO1*) also resulted in higher sensitivity to 5hm-dC, which was not replicated in the genome-wide screen. This discrepancy might be explained by the fact that DT40 cells lack *TRP53* (21), resulting in a lack of G1/S phase checkpoint activation in response to damage, and increased susceptibility to DNA re-replication (22). Therefore, DT40 MMR mutants may be disproportionally sensitive to exogenous genomic threats, including 5hm-dC if they induce damage during G1 and/or S phase, due to the requirement for MMR in S/G2 phase checkpoint activation (23, 24).

Finally, to confirm the involvement of BER in 32D cells, we generated a CRISPR/Cas9 knockout cell line (**Supplementary Fig. 1D**) and measured growth inhibition in response to different concentrations of 5hm-dC using a fluorescent or colorimetric cell viability assay. We found that *Polb^−/−^* 32D cells were hypersensitive to 5hm-dC treatment compared to a wildtype control (**Fig. 1E**). This uncovers the need for BER and specifically POLβ to protect cells from 5hm-dC mediated sensitivity. Collectively, these findings indicate that multiple DNA repair pathways contribute to cellular tolerance of exogenous 5hm-dC, with BER playing a predominant role, and demonstrate that the requirement for BER and specifically POLβ is conserved across vertebrate species.

### Base excision repair protects cells from genotoxicity by exogenous 5hm-dC

As 5hmC is a naturally occurring modified base, we hypothesized that intracellular accumulation and its incorporation into DNA may result in DNA damage necessitating BER-mediated resolution. To test this, we utilized immunofluorescence to visualise RPA foci (a marker of SSBs) and γH2A.X (a marker of DSBs). After exposing cells to 5hm-dC, we observed that while wildtype cells accumulate a significant increase of RPA and γH2AX foci at 200 μM, *Polb^−/−^* cells display a 21-fold and 52-fold higher increase, respectively (**Fig. 2A-B**) Overall, this shows that BER is required to prevent the formation of DNA breaks when exposed to 5hm-dC.

**Figure 2.**
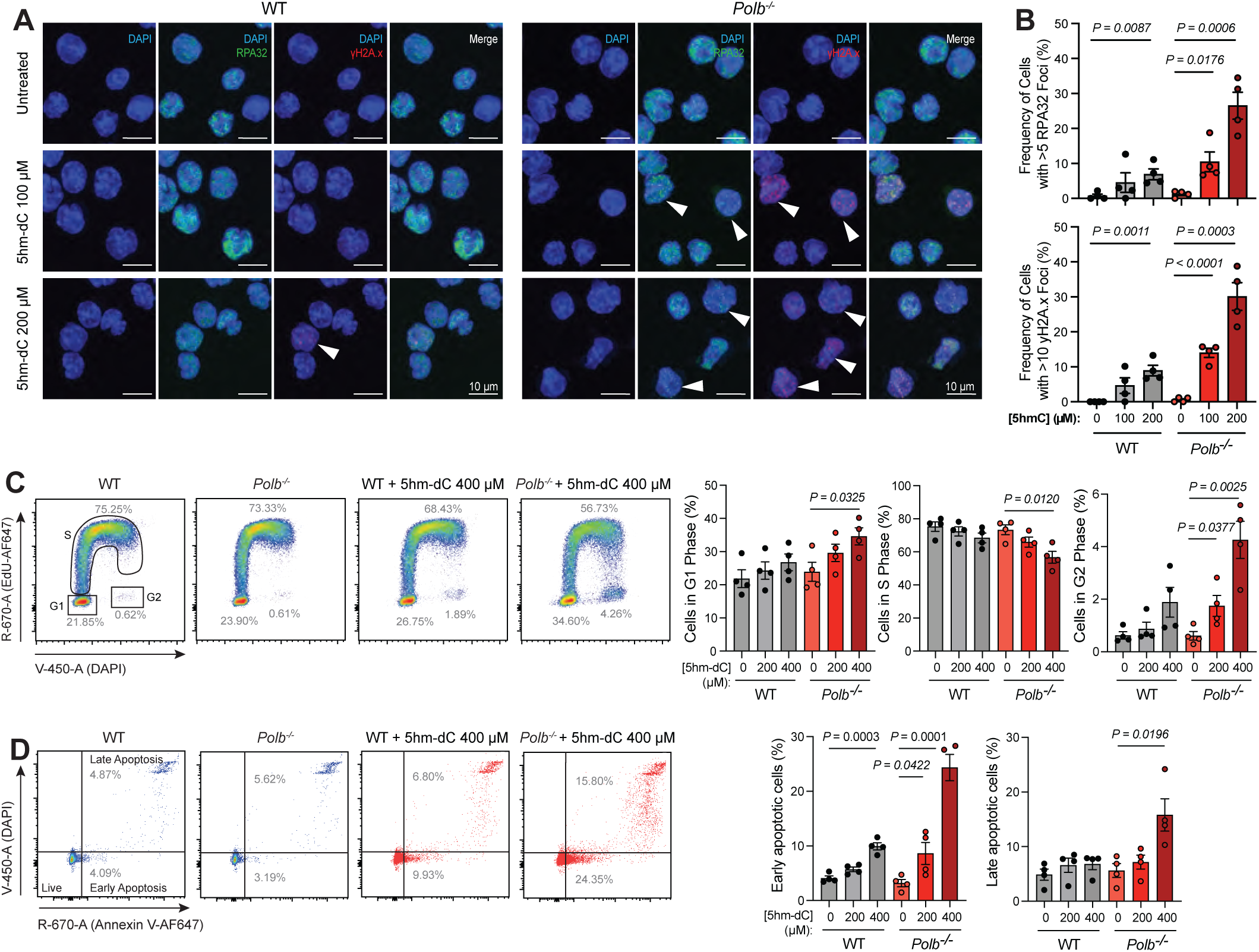
Exogenous 5hm-dC induces DNA damage, cell cycle arrest, and apoptosis in BER deficient *Polb^−/−^*cells. **A)** Representative images of gH2A.X and RPA32 foci in WT and *Polb^−/−^*32D cells in response to increasing 5hm-dC concentrations. Arrows indicate cells with >5 RPA32 or >10 gH2A.X foci. Scale bar, 10µM. **B)** Quantification of the percentage of WT and *Polb^−/−^*32D cells with >5 RPA32 foci (left) and >10 gH2A.X foci (right). Data represents the mean and s.e.m. of n=4 biological replicates. **C)** Representative flow cytometry plots of EdU cell cycle analysis in WT and *Polb^−/−^* 32D cells in response to increasing 5hm-dC concentrations. Percentage values represent the mean across experimental replicates and quantification of the percentage of cells in G1, S and G2 phases for WT and *Polb^−/−^* 32D cells in response to indicated 5hm-dC concentrations. Data represents the mean and s.e.m. of n=4 biological replicates. **D)** Representative flow cytometry plots of early and late apoptotic cell populations in WT and *Polb^−/−^* 32D cells in response to increasing 5hm-dC concentrations. Percentage values represent the mean across experimental replicates. Quantification of the percentage of cells in early (Annexin V -positive, DAPI-negative) and late (Annexin V -positive, DAPI-positive) apoptosis for WT and *Polb^−/−^*32D cells in response to indicated 5hm-dC concentrations. Data represents the mean and s.e.m. of n=4 biological replicates. *P* values were calculated using an unpaired t-test (only *P* values < 0.05 are displayed).

The accumulation of DNA damage leads to cell cycle arrest and, ultimately, cell death. To explore whether exogenous 5hm-dC treatment alters cell cycle dynamics in wildtype and *Polb^−/−^* cells, we analysed the cell cycle using flow cytometry utilizing the analogue ethynyl-2’-deoxyuridine (EdU), actively incorporated during the S phase, in combination with a DNA dye. After a 72-hour incubation with 200 or 400 μM 5hm-dC, wildtype cells showed no significant difference in the proportion of cells in each phase with or without treatment (**Fig. 2C**). However, in *Polb^−/−^* cells we observed a significant increase in the proportion of cells in the G1 and G2 phases, and a concomitant decrease in S phase cells. This shows that in the absence of BER, 5hm-dC-mediated DNA damage leads to cell cycle arrest in either G1 or G2. This may reflect triggering of the G1/S and G2/M damage-induced cell cycle checkpoints.

Finally, we determined whether BER deficiency leads to apoptotic cell death following 5hm-dC exposure. Wildtype and *Polb^−/−^* cells were treated with 200 or 400 µM 5hm-dC for 72 h and analysed by Annexin V FACS. Wildtype cells showed increased early apoptosis only at 400 µM, with no change in late apoptosis (**Fig. 2D**). In contrast, *Polb^−/−^* cells exhibited elevated early apoptosis at both concentrations and a significant increase in late apoptosis at 400 µM. These data indicate that loss of BER enhances apoptotic sensitivity to exogenous 5hm-dC, consistent with the reduced viability observed in *Polb^−/−^* cells (**Fig. 1E**). In summary, these results show that BER is essential to protect cells from cell cycle arrest, DNA damage and eventually cell apoptosis when exposed to exogenous 5hm-dC.

### Genotoxicity by exogenous 5hm-dC arises due to metabolic conversion into 5hm-dU

The requirement of BER for toleration of exogenous 5hm-dC suggests that exogenous 5hm-dC mediates some form of base damage. To determine the molecular mechanism of cellular toxicity we utilized an unbiased genome-wide CRISPR/Cas9 screen in *Polb^−/−^* 32D cells to identify factors whose loss allows BER-deficient cells to survive in the presence of exogenous 5hm-dC.

The screening was performed as described above, now using Cas9-expressing *Polb^−/−^* cells, generating a pool of double knockout cells (**Fig. 3A**). This time the hits of interest were the guides that are enriched after 5hm-dC treatment, the resistant population. Using the MAGeCK analysis described previously, we can identify positively selected genes in a screen. Among the top hits we consistently found genes involved in nucleotide metabolism (*Slc29a2, Dck, Dctd),* BER (*Smug1*) and transcriptional regulation (*Cir1*, *Fbxo42*) (**Fig. 3B, Supplementary Fig. 2**).

**Figure 3.**
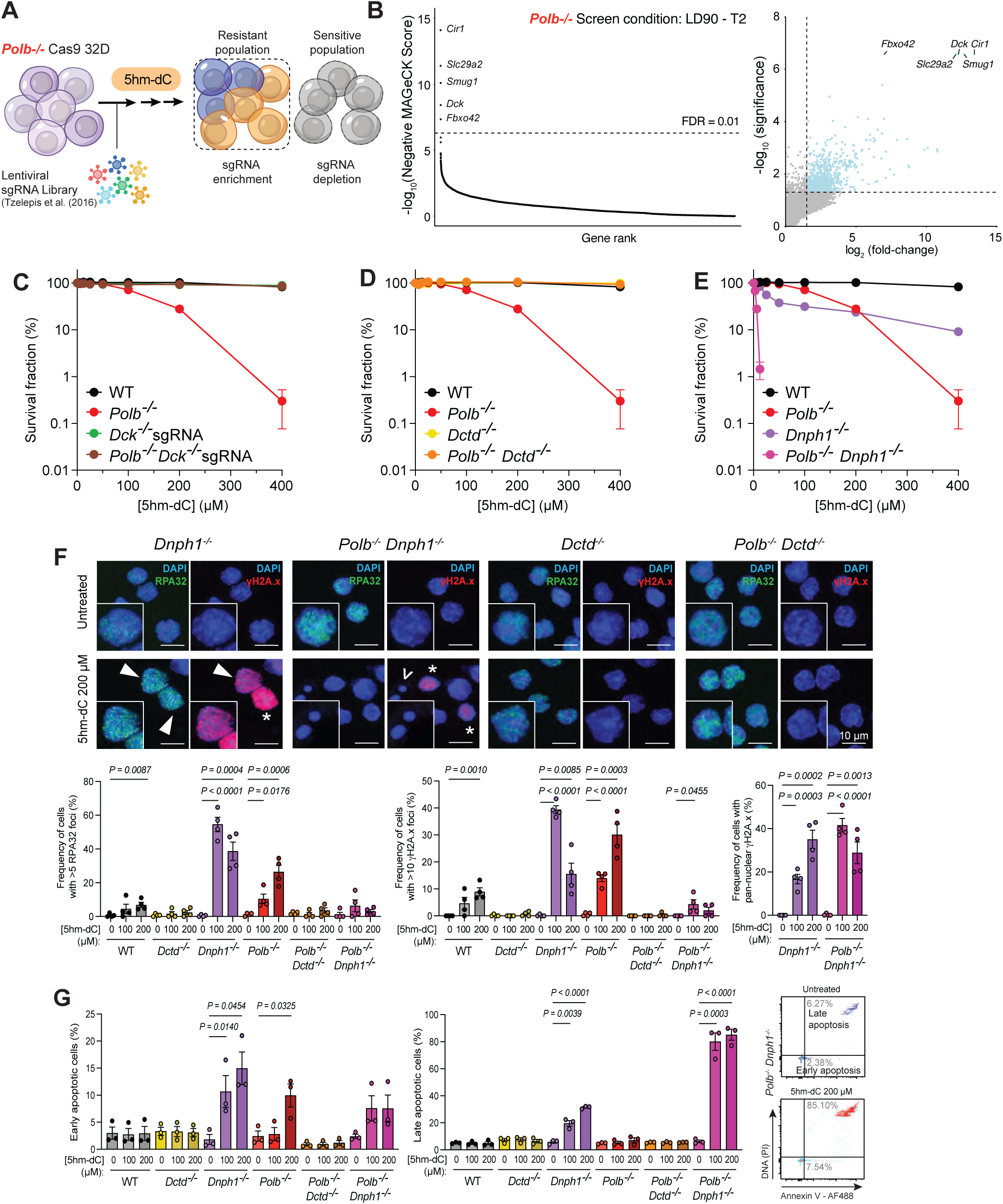
Genome wide CRISPR suppressor screen identifies dNTP metabolism as key determinate of 5hm-dC toxicity in *Polb^−/−^* cells. **A)** Schematic overview of the genome-wide CRISPR/Cas9 sensitivity screen in 32D mouse myeloid progenitor cells. Cas9-expressing 32D wildtype and *Polb^−/−^* cells were transduced with a whole-genome sgRNA library and exposed to 5-hydroxymethyl-2′-deoxycytidine (5hm-dC). Samples were collected prior to treatment (T0) and after two and three sequential treatments (T1 and T2). sgRNA representation was determined by next-generation sequencing to identify depleted and enriched guides. **B)** MAGeCK analysis of sgRNA abundance comparing untreated and 5hm-dC-treated cells under LD90 at T2. Genes were ranked based on statistical significance, fold change, MAGeCK score and false discovery rate. The left column shows screen results ranked by MAGeCK score, indicating the FDR cutoff (FDR < 0.1). The right column shows volcano plots with the significance scores from MAGeCK analysis. The top five ranked hits according to MAGeCK score for each condition are labelled. **C-E)** Survival curves of WT, *Polb^−/−^* and the indicated single-double-knockout cell lines (*Dck^−/−^* sgRNA-targeted, *Dctd^−/−^* clonal and *Dnph1^−/−^* sgRNA-targeted) in response to increasing 5hm-dC concentration. Data represents the mean and s.e.m. of n=3 biological repeats (n=3 technical repeats each) **F)** Representative images of gH2A.X and RPA32 foci for indicated cell lines. Arrows indicate cells with >5 RPA32 or >10 gH2A.X foci, asterisks represent cells with pan-nuclear gH2A.X, hollow arrows represent apoptotic cells. Scale bar, 10µM. Corresponding quantification of the percentage of >5 RPA32 foci (left) and >10 gH2A.X foci (middle) pan-nuclear gH2A.X (left). Data represents the mean and s.e.m. of n=4 biological replicates. **G)** Quantification of the percentage of cells in early (Annexin V-positive, PI-negative) and late (Annexin V-positive, PI-positive) apoptosis for indicated cell lines. Data represents the mean and s.e.m. of n=3 biological replicates. *P* values were calculated using an unpaired t-test (only *P* values < 0.05 are displayed). Representative flow cytometry plots are shown on the right.

We first turned our attention to nucleotide metabolism. *Slc29a2* encodes Equilibrative Nucleoside Transporter-2 (ENT2), a bidirectional uniporter involved in nucleoside transport. This protein confirmed the effectiveness of our screen, as it is necessary for import of 5hm-dC into the cell, as inhibition using ENTi results in reduced sensitivity of *Polb^−/−^* to 5hm-dC (**Supplementary Fig. 3A**). Next, we find *Dck* as one of the other top hits (**Fig. 3B, Supplementary Fig. 2**). DCK catalyses the phosphorylation of deoxycytidine and analogues to generate deoxycytidine-5’-monophosphate (dCMP), the first step in generating a deoxynucleotide-triphosphate (dNTP). We hypothesized that DCK must also act on 5hm-dC, as a *Polb^−/−^ Dck^−/−^* double knockout does not display a growth disadvantage to wildtype cells upon treatment (**Fig 3C, Supplementary Fig. 3**). Thus, 5hm-dC-mediated genotoxicity is dependent on its phosphorylation to 5hm-dCMP and incorporation into the nucleotide pool. Interestingly, we also find *Dctd* in the suppressor screen in both the LD50 and LD90 T1 condition, as the 10th and 12th ranked hits respectively (**Supplementary Source data**). DCTD is another enzyme involved in pyrimidine metabolism, normally responsible for the deamination of dCMP to dUMP to synthesize dTMP (25). We therefore speculated that DCTD might be responsible for the deamination of 5hm-dCMP to 5hm-dUMP in the nucleotide pool. We knocked out the *Dctd* gene and found that *Polb^−/−^* cells also lacking *Dctd* are now completely resistant to 5hm-dC (**Fig. 3D, Supplementary Fig. 3**). These results indicate that DCTD-mediated deamination in the nucleotide pool is responsible for the genotoxicity caused by 5hm-dC.

In earlier sensitivity screening, we also observed that the *Dnph1* gene was one of the highest-ranking hits (**Fig. 1B**). This gene has recently been shown to be involved in sanitizing the nucleotide pool from 5hm-dUMP (15). We found that when we knockout out *Dnph1*, both *Dnph1^−/−^* and *Dnph1^−/−^Polb^−/−^* cells are hyper-sensitive to 5hm-dC treatment (**Fig. 3E, Supplementary Fig. 3D**). Most notably, the double knockout of *Dnph1* and *Polb* leads to a significant increase in the 5hm-dC sensitivity, indicating that an increased 5hm-dUMP pool severely exacerbates the phenotype. These results argue that 5hm-dCMP is deaminated in the nucleotide pool but alternatively, we considered the possibility that 5hm-dC may be incorporated and deaminated on DNA. We find that this is unlikely to be the case as knockout of *Aicda*, *Apobec1*, *Apobec2*, and *Apobec3*, the proteins that deaminate bases on the DNA, do not result in rescue of *Polb^−/−^* hypersensitivity (**Supplementary Fig. 4A-B**), suggesting that it is instead deaminated in the nucleotide pool.

Focusing on the deamination of the nucleotide pool, we further tested if the loss of *Dctd* or *Dnph1* would rescue or exacerbate genotoxicity respectively by measuring both DNA damage and cell apoptosis as described before. We found that cells that lose *Dctd* have their DNA damage and late and early apoptosis levels reduced back to the same extent as wildtype cells (**Fig. 3F-G, Supplementary Fig. 5A-B**). For *Dnph1*, we see a significant increase in both RPA and γH2AX foci in the single knockout, however at higher concentration foci seem to go down but accompanied by an increased in pan-nuclear γH2AX staining (**Fig. 3F, Supplementary Fig. 5A**). In the *Dnph1^−/−^ Polb^−/−^* cells there does not seem to be a large increase in RPA and γH2AX foci, however here there is a larger increase in pan-nuclear staining of γH2AX. In agreement with this, most cells are in a late apoptotic state (**Fig. 3G, Supplementary Fig. 5B**), unlike *Dnph1^−/−^* cells, as indicated by the accumulation of small, spherical DAPI-stained apoptotic bodies (**Supplementary Fig. 5A**). Together, the data presented here delineate a mechanism where 5hm-dC enters the cell through ENT2 and the nucleotide pool following phosphorylation by DCK. 5hm-dCMP is likely deaminated in the nucleotide pool by DCTD and 5hm-dUMP which requires DNPH1 for nucleotide pool sanitation.

### Excision of 5hmU from DNA drives genotoxicity to exogenous 5hm-dC

An important hit from the screen was the DNA glycosylase *Smug1* across multiple conditions (**Fig. 3B, Supplementary Fig. 2**). SMUG1 is involved in excising uracil and uracil analogues from the DNA, including 5hmU, a deaminated version of 5hmC, but excluding cytosine bases (8, 26). In agreement with previous reports (16), we see that treatment of POLB-deficient cells with exogenous 5hm-dU also results in genotoxicity (**Supplementary Fig. 6**). This suggests that DNA lesions caused by exogenous 5hm-dC treatment may instead be a consequence of SMUG1-mediated 5hmU excision, and the inability to repair the resulting BER intermediate.

To validate this observation, we knocked out *Smug1* using lentiviral transduction with a *Smug1*-targeting sgRNA, in both wildtype and *Polb^−/−^* 32D cells (**Supplementary Fig. 6A-C**). When exposing these cells and measuring growth inhibition we found that hypersensitivity found in *Polb^−/−^* cells is completely lost upon knockout of *Smug1* (**Fig. 4A**). This suggests that sensitivity is caused by the removal of 5hmU during the upstream process in BER.

**Figure 4.**
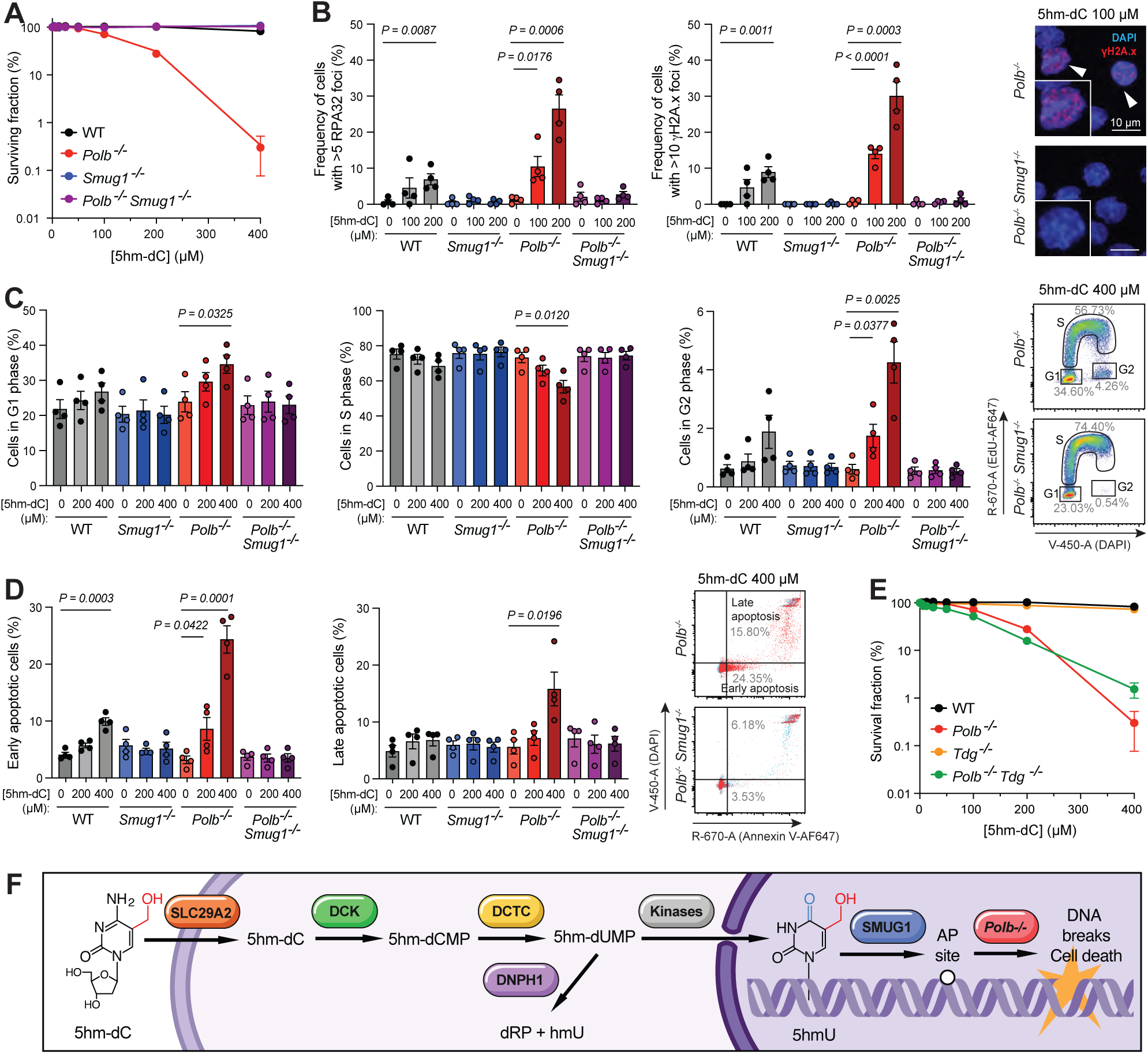
*Smug1* mediated excision of 5hmU drives 5hm-dC induced genotoxicity in *Polb^−/−^* cells. **A)** Survival curves of WT, *Polb^−/−^*, *Smug1^−/−^* sgRNA-targeted pool, and *Polb^−/−^ Smug1^−/−^* sgRNA-targeted pool cell lines in response to increasing 5hm-dC concentration. Data represents the mean and s.e.m. of n=3 biological repeats (n=3 technical repeats each). **B)** Quantification of the percentage of cells with >5 RPA32 foci (left) and >10 gH2A.X foci (right) in the indicated cell lines. Data represents the mean and s.e.m. of n=4 biological replicates. On the right, representative images with gH2A.X foci. **C)** Quantification of the percentage of cells in G1 (left), S (middle) and G2 (right) phases for indicated cell lines in response to 5hm-dC. Data represents the mean and s.e.m. of n=4 biological replicates. On the right, representative flow cytometry plots. **D)** Representative flow cytometry plots of early and late apoptosis in the indicated cell lines following 5hm-dC treatment, with corresponding quantification of early (Annexin V-positive, DAPI-negative) and late (Annexin V-positive, DAPI-positive) apoptosis. Data represents the mean and s.e.m. of n=4 biological replicates. On the right, representative flow cytometry plots. **B-D)** *P* values were calculated using an unpaired t-test (only *P* values < 0.05 are displayed). **E)** Survival curves of WT, *Polb^−/−^*, *Tdg^−/−^*, and *Polb^−/−^ Tdg^−/−^* clonal cell lines in response to increasing 5hm-dC concentration. Data represents the mean and s.e.m. of n=3 biological repeats (n=3 technical repeats each). **F)** Model for the toxicity of 5hm-dC. 5hm-dC enters the cell via the SLC29A2 nucleoside transporter. Once in the cytoplasm, *DCK* phosphorylates the nucleoside to generate 5hmdCMP. *DCTD* deaminates 5hmdCMP to generate 5hmdUMP. 5hmdUMP is subject to two routes of further processing. The DNHP1 hydrolase may sanitise 5hmdUMP from the nucleotide pool by cleaving the moiety to generate 5hmU and dRP (deoxyribosephosphate). Alternatively, 5hmdUMP may be subject to further phosphorylation by cytoplasmic kinases, generating 5hmdUTP that is misincorporated into genomic DNA. Genomic 5hmU is excised by the *SMUG1* DNA glycosylase, initiating BER to repair the resulting abasic site. Excessive genomic incorporation of 5hm-dUTP may cause genotoxicity by overloading the BER capacity of the cell, such that *SMUG1*-mediated abasic sites cannot be repaired, leading to genotoxicity and cell death.

Next, we sought to confirm if SMUG1-mediated base excision also underlies the DNA damage, cell cycle arrest and apoptosis in *Polb^−/−^* cells. In both *Smug1^−/−^* and *Smug1^−/−^ Polb^−/−^* cells, we observe that exogenous 5hm-dC exposure does not induce accumulation of RPA or γH2AX foci (**Fig. 4B, Supplementary Fig. 6D**). This indicates that in the absence of SMUG1 activity, cells fail to obtain SSBs nor DSBs. We also observe that lack of *Smug1* completely rescues the cell cycle defects observed in *Polb^−/−^* cells with no accumulation of cells in G1 or G2 phase (**Fig. 4C, Supplementary Fig. 6E**). Lastly, knockout or *Smug1* also results in complete rescue of the cell apoptosis phenotype found in *Polb^−/−^* cells at all concentrations (**Fig. 4D, Supplementary Fig. 6F**).

On DNA, 5hmC is also further oxidized to 5fC and 5caC by the TET proteins. Considering the role of SMUG1 mediated removal of 5hmU, we questioned the involvement of the oxidized bases 5fC and 5caC. Canonically, these bases are removed during active DNA demethylation by the DNA glycosylase TDG in a BER dependent manner. To this extent we generated *Tdg^−/−^* and *Tdg^−/−^ Polb^−/−^* 32D cells (**Supplementary Fig. 7A**) and measured cell viability using a survival assay after exposure to 5hm-dC. We found that unlike *Smug1*, knocking out *Tdg* in *Polb^−/−^* cells did not affect the hypersensitivity (**Fig. 4E**), generation of DNA breaks, apoptosis and had a minimal effect on cell cycle (**Supplementary Fig. 7B-D**). This indicates that TDG does not play a major role in exogenous 5hm-dC-induced damage.

We propose a model for exogenous 5hm-dC genotoxicity (**Fig. 4F**), in which 5hm-dC is imported into 32D cells via SLC29A2, phosphorylated by DCK to 5hm-dCMP, and deaminated by DCTD to 5hm-dUMP. DNPH1 normally sanitises the nucleotide pool by converting 5hm-dUMP to 5hmU, preventing formation of 5hm-dUTP. Without DNPH1, 5hm-dUTP accumulates and is misincorporated into DNA as 5hmU. SMUG1 excises 5hmU via BER, generating abasic sites. In *Polb^−/−^* cells, unresolved BER intermediates accumulate as SSBs and DSBs, triggering cell cycle arrest and apoptosis.

### Base excision repair is essential to protect cells during active DNA demethylation

We used exposure to the nucleoside 5hm-dC as an experimental system to understand genotoxicity by 5hmC and found that it is actually mediated by 5hmU. Next, we wanted to ask whether endogenous generation of 5hmC during active DNA demethylation is similarly genotoxic, and whether oxidative demethylation is a significant source of 5hmU as proposed by others (15). To study this, we generated an inducible system where a doxycycline-inducible (Tet-On) promotor drives expression of the catalytic domain of TET1 (**Fig. 5A**), knocked into the TIGRE locus (27). This construct includes an EGFP protein following a T2A peptide allowing fluorescence to be used as a proxy for TET1 expression (**Fig. 5A, Supplementary Fig. 8A-C**). Two versions of the construct were used, one containing the wild type catalytic domain of TET1 (TET1 CD^WT^) and a catalytic-dead H1672Y/D1674A point mutant (TET1 CD^PM^) (28). After 24 hours treatment with doxycycline, we could readily detect EGFP expression by FACS (**Fig. 5A**). But to confirm that this system also induces active DNA demethylation, we used LC-MS to quantify 5mC and its oxidation products.

**Figure 5.**
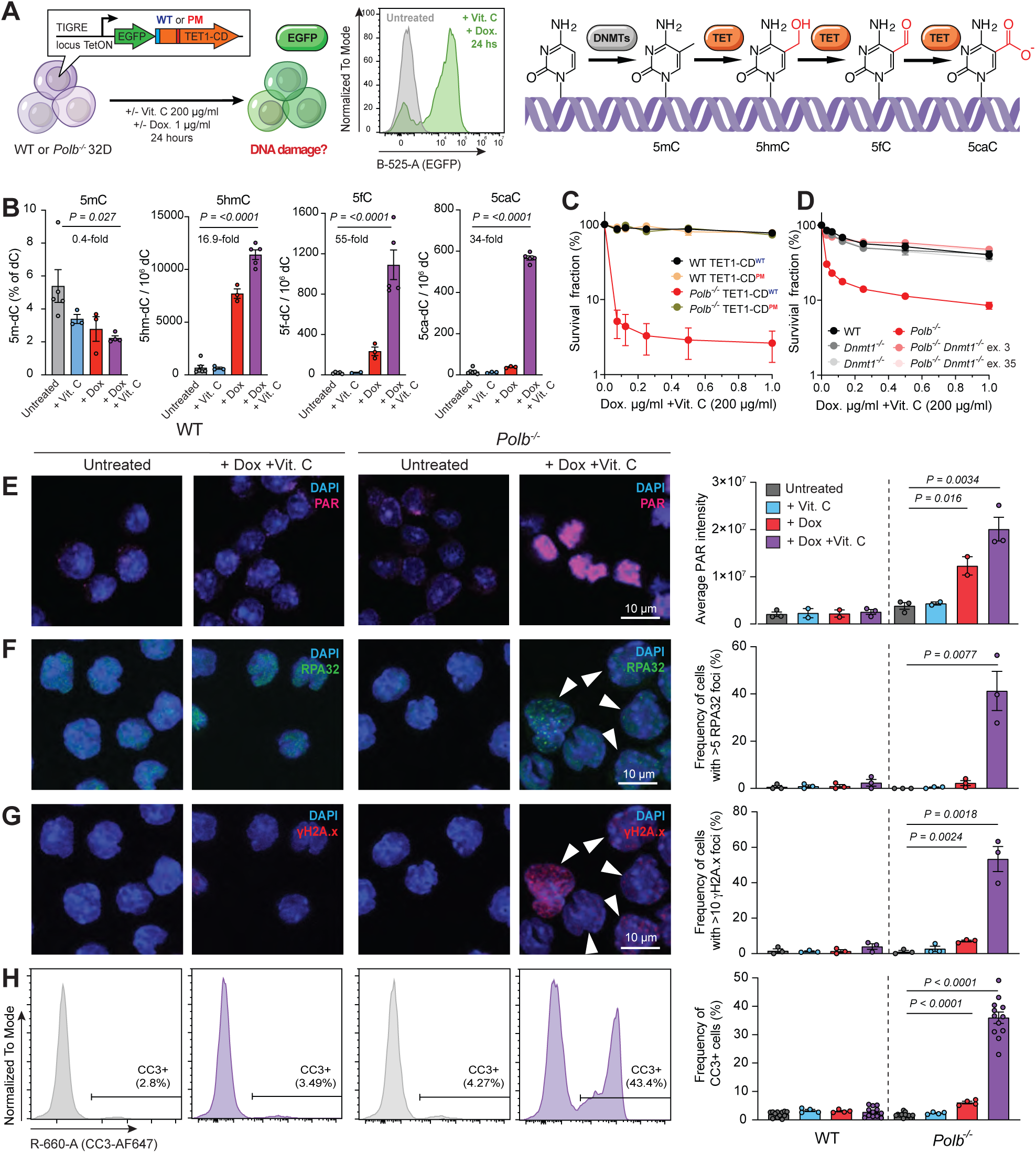
Endogenous TET1 mediated oxidative DNA demethylation induces BER-dependent genotoxcity in *Polb^−/−^* cells. **A)** Schematic model of the doxycycline-inducible tet-on TET1-CD^wt^/TET1-^CDH1672YD1674A(PM)^ system, with representative flow cytometry plots demonstrating EGFP expression following induction with 1µg/mL doxycycline. **B)** Quantification of DNA methylation and oxidation products by LC-MS in WT TET1-CD^wt^ in response to 1µg/mL doxycycline and 200µg/mL Vitamin C. Data represents the mean and s.e.m. of n=3 biological repeats **C)** Survival curves of WT TET1-CD^wt^/TET1-CD^PM^, and *Polb^−/−^* TET1-CD^WT^/TET1-CD^pm^ clonal cell lines in response to increasing doxycycline concentration and 200µg/mL Vitamin C. Data represents the mean and s.e.m. of n=3 biological repeats (n=3 technical repeats each). **D)** Survival curves of WT, *Dnmt1^−/−^*, *Polb^−/−^*, *Polb^−/−^ Dnmt1^−/−^*, sgRNA-targeted pool containing the TET1-CD^wt^ construct, in response to increasing doxycycline concentration and 200µg/mL Vitamin C. Data represent the mean and s.e.m. of n=3 biological repeats (n=3 technical repeats each). **E-G)** Representative images of Parylation (PAR)(E) RPA32 (F) and gH2A.X (G) foci in WT TET1-CD^WT^ and *Polb^−/−^* TET1-CD^WT^ cells following induction with 1µg/mL doxycycline and 200µg/mL Vitamin C. Arrows indicate cells with >5 RPA32 or >10 gH2A.X foci. Scale bar, 10µM. Corresponding quantification is shown. Data represents the mean and s.e.m. of n=3 biological replicates (PAR: n = 3 for untreated and combined treatment, n = 2 for single treatments) **H)** Representative flow cytometry plots of apoptotic cell populations, by cleaved caspase 3, in WT TET1-CD^WT^ and *Polb^−/−^* TET1-CD^WT^ clonal cell lines following induction with 1µg/mL doxycycline and 200µg/mL Vitamin C. Percentage values represent the mean across experimental replicates. Corresponding quantification of the percentage of cell apoptosis. Data represents the mean and s.e.m. of n = 12 biological replicates (untreated and combined treatment) and n = 4 (single treatments). *P* values were calculated using an unpaired t-test (only *P* values < 0.05 are displayed).

The quantification was performed using isotope dilution liquid chromatography tandem mass spectrometry (LC-ID-MS/MS), following an extended version of a methodology previously reported by some of us (29), with the addition of 5hm-dU and 5m-dC. Briefly, DNA was isolated, purified, and enzymatically digested prior to spiking with isotopically labelled internal standards (**Supplementary Fig. 9A**). The resulting digest was subjected to HPLC analysis to enable the parallel quantification of unmodified nucleosides and enrichment of epigenetic modifications which were subsequently analysed by LC-ID-MS/MS (**Supplementary Fig. 9B-E, Methods**).

Initial experiments with doxycycline alone showed modest increases in oxidative demethylation (**Fig. 5B**). We therefore added vitamin C as a cofactor to further enhance DNA demethylation, as it maintains catalytic Fe²⁺ levels and thereby promotes iterative oxidation of 5mC to 5hmC, 5fC and 5caC (30). We detected approximately 5% of cytosines to be methylated, and double treatment resulted in an approximately 40% reduction in total genomic 5mdC levels and a marked increase in all oxidation products, with 5hm-dC, 5f-dC and 5ca-dC levels increasing by approximately 17-, 55- and 38-fold relative to total dC, respectively. (**Fig. 5B**). These levels are comparable to those observed in neurons (31–33), suggesting our system recapitulates physiologically relevant demethylation levels. This data confirms that our inducible system effectively induces oxidative DNA demethylation.

Next, we assessed whether BER and POLβ are required to protect cells from TET1-induced demethylation. Using a growth inhibition assay we observed that wildtype cells are unaffected at lower concentrations of doxycycline, independent of the catalytic activity of TET1, as no difference is seen between the TET1-CD^WT^ and TET1-CD^PM^. However, *Polb^−/−^* cells are hypersensitive to the overexpression of TET1-CD^WT^ but not catalytic-dead TET1-CD^PM^ (**Fig. 5C**). Interestingly, the toxicity by TET1 CD^WT^ required the expression of DNMT1, responsible for the maintenance of 5mC, indicating that cytosine methylation was necessary for this effect and it is not a secondary effect from the overexpression TET1 proteins (**Fig. 5D**).

We previously demonstrated that exogenous 5hm-dC causes DNA damage. Similarly, we found that *Polb^−/−^* cells have increased PARylation (another marker of SSBs), RPA32 foci and yH2AX foci (**Fig. 5E-G, Supplementary Fig. 10**) compared to wild type cells, after expression of TET1 CD^WT^, but not TET1 CD^PM^. Furthermore, we see that while wild type cells do not have an increase in apoptosis measured by either cleaved caspase 3 (CC3) or Annexin V (**Fig. 5H, Supplementary Fig. 11),** *Polb^−/−^* cells show increased cells dead. Together, these data demonstrate that TET1-mediated oxidative DNA demethylation generates base lesions that require BER-mediated processing to maintain cellular viability, and that failure to resolve these lesions in the absence of POLβ leads to DNA damage accumulation and cell death.

### Acute genotoxicity from DNA demethylation in POLβ-deficient cells is mediated by TDG

As exogenously supplied 5hm-dC induces sensitivity in BER-deficient cells through deamination in the nucleotide pool to 5hm-dU, we next asked whether a similar mechanism operates during endogenous TET-mediated DNA demethylation. Specifically, we sought to determine whether genotoxicity arises via the same mechanism, or through another pathway, as oxidative demethylation has also been proposed as a potential source of 5hmU (15).

To explore this, we generated *Tdg^−/−^*, *Smug1^−/−^* and single or *Polb^−/−^* double knockout cells containing the TET1 CD^WT^ or TET1 CD^PM^ constructs (**Supplementary Fig. 12A-D**). We first assessed DNA damage markers by RPA or yH2AX foci, 12 hours after induction of oxidative demethylation. Strikingly, loss of *Tdg*, but not *Smug1*, completely abolished the formation of both SSBs and DSBs in the *Polb^−/−^* background (**Fig. 6A-C, Supplementary Fig. 12E**). In addition, *Tdg* knockout completely suppressed apoptosis, whereas loss of either *Smug1* or *Dnph1* did not alter cell death compared with their respective wild-type or *Polb^−/−^* controls, as measured by CC3 and Annexin V staining at 24 hours (**Fig. 6D, Supplementary Fig. 13**). This contrasts with exogenous 5hm-dC treatment, where *Smug1* or *Dnph1* inactivation suppressed or exacerbated the *Polb^−/−^* phenotype, respectively; we did not observe the same effects following TET1 expression. These results indicate that deamination of 5hm-dC in the nucleotide pool or deamination of 5hmC on DNA is unlikely to contribute to genome instability immediately after the acute induction of endogenous DNA demethylation. Instead, oxidation to 5f-dC and 5ca-dC, followed by their excision and AP site formation is likely responsible.

**Figure 6.**
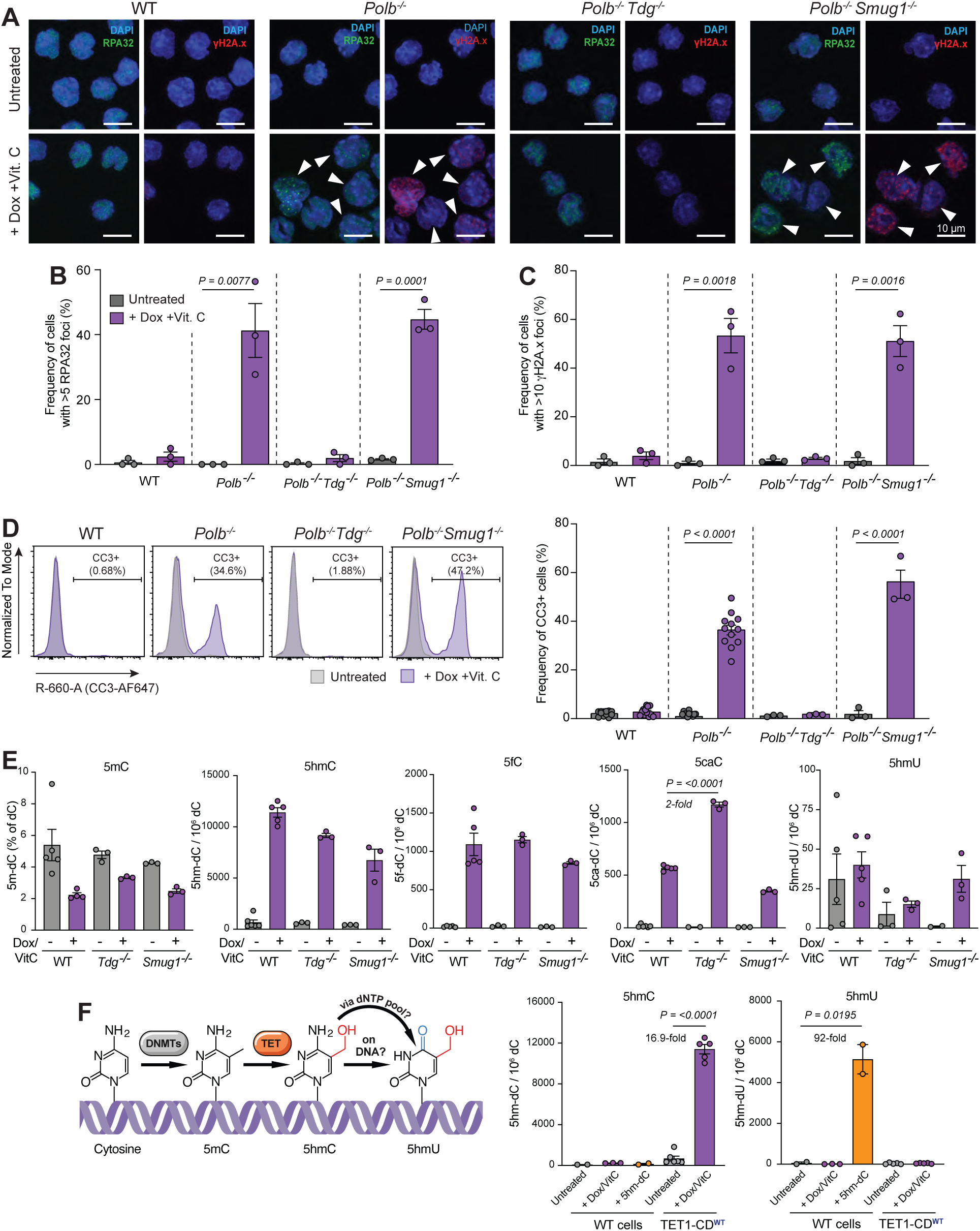
Oxidative DNA demethylation induced genotoxicity is mediated by *Tdg*-dependent base excision rather than 5hmU processing. **A-C)** Representative images of RPA32 and gH2A.X foci (A) and quantifications of cells with >5 RPA32 (B) or >10 gH2A.X (C) foci in WT-, *Polb^−/−^*-, *Polb^−/−^ Tdg^−/−^*-, *Polb^−/−^ Smug1^−/−^*-TET1-CD^WT^ clonal cell lines following induction with 1µg/mL doxycycline and 200µg/mL Vitamin C. Arrows indicate cells with >5 RPA32 foci and >10 gH2A.X foci. Scale bar, 10µM. Data represents the mean and s.e.m. of n=3 biological replicates. **D)** Representative flow cytometry plots of apoptotic cell populations, by cleaved caspase 3, in WT-, *Polb^−/−^*-, *Polb^−/−^ Tdg^−/−^*-, *Polb^−/−^ Smug1^−/−^*-TET1-CD^WT^ clonal cell lines following induction with 1µg/mL doxycycline and 200µg/mL Vitamin C. Percentage values represent the mean across experimental replicates. Corresponding quantification of the percentage of cells apoptosis. Data represents the mean and s.e.m. of n=12 biological replicates for WT- and *Polb^−/−^*-, n=3 for *Polb^−/−^ Tdg^−/−^*- and *Polb^−/−^ Smug1^−/−^*-TET1-CD^WT^. *P* values were calculated using an unpaired t-test (only *P* values < 0.05 are displayed). **E)** Quantification of DNA methylation and oxidation products by LC-MS in WT-, *Tdg^−/−^*-, *Smug1^−/−^*-TET1-CD^wt^ in response to 1µg/mL doxycycline and 200µg/mL Vitamin C. Data represents the mean and s.e.m. of n=3 biological repeats. **F)** Quantification and direct comparison of 5hmC and 5hmU levels in response to exogenous 5hm-dC treatment (750 µM) or oxidative demethylation induced 1µg/mL doxycycline and 200µg/mL Vitamin C treatment. Data represents the mean and s.e.m. of n = 2 biological replicates (untreated and 5hm-dC WT cells), n = 3 (1µg/mL doxycycline and 200µg/mL Vitamin C treatment WT cells), n = 5 (1µg/mL doxycycline and 200µg/mL Vitamin C treatment TET1-CD^WT^ cells) and n = 6 (untreated TET1-CD^WT^ cells) biological replicates.

To further validate this genetic finding, we aimed to directly quantify epigenetic bases in the *Tdg^−/−^* and *Smug1^−/−^* excision-deficient mutants using LC-ID-MS/MS. Following 24 hours of doxycycline and vitamin C treatment, all cell lines showed decreased levels of 5mC and corresponding increase in the levels of 5hmC indicative of similar TET1 activity (**Fig. 6E**). As expected, *Tdg^−/−^* cells accumulated higher levels of 5f-dC and 5ca-dC compared to wild type cells, in agreement with the known role of TDG in excising these modified bases. Finally, we detected very variable levels of 5hmU and no obvious induction linked to oxidative demethylation, despite the 17-fold increase 5hmC, suggesting 5hmC is not readily deaminated on DNA. This result contrasts with the 92-fold increase in 5hmU following exogenous 5hm-dC exposure in wild-type cells (**Fig. 6F**). Together, these data demonstrate that TET1-mediated DNA damage and cell death POLβ-deficient cells is caused by TDG-mediated excision of 5f-dC and 5ca-dC, and that oxidative demethylation is not a major source of 5hmU.

### Oxidative DNA demethylation in cytotoxic in the absence of SMUG1 or DNPH1

Finally, we examined the effects of prolonged TET1 activation over a longer period. We reasoned that if oxidative demethylation was a significant source of 5hm-dU, 24 hours may not be sufficient for 5mC to be oxidised, excised, deaminated in the nucleotide pool and incorporated back into DNA during replication. First, we exposed cells to doxycycline with vitamin C in a growth inhibition assay continuously for 72 hours. We found that knockout of *Tdg* completely rescued the phenotype of *Polb^−/−^* cells (**Fig. 7A**), in agreement with the suppression of DNA damage and apoptosis (**Fig. 6A-D**). Surprisingly, we found that *Smug1^−/−^* and *Dnph1^−/−^* single mutants, and their *Polb^−/−^* double mutants, are more sensitive than their respective controls (**Fig. 7A**). The decreased survival was detected in both the single and *Polb^−/−^* double knockouts, suggesting that the toxicity occurs independently of BER. At first glance, this was inconsistent with our previous measurements of apoptosis at 24 hours (**Fig. 6D, Supplementary Fig. 13**). Therefore, we repeated this experiment but measured apoptosis at different time points before and during a longer doxycycline and vitamin C treatment. This time course analysis revealed an induction of CC3+ apoptotic cells in *Smug1^−/−^*, *Dnph1^−/−^*, and their *Polb^−/−^* double mutants at 48 and 72 hours (**Fig. 7B**), in agreement with the growth inhibition data.

**Figure 7.**
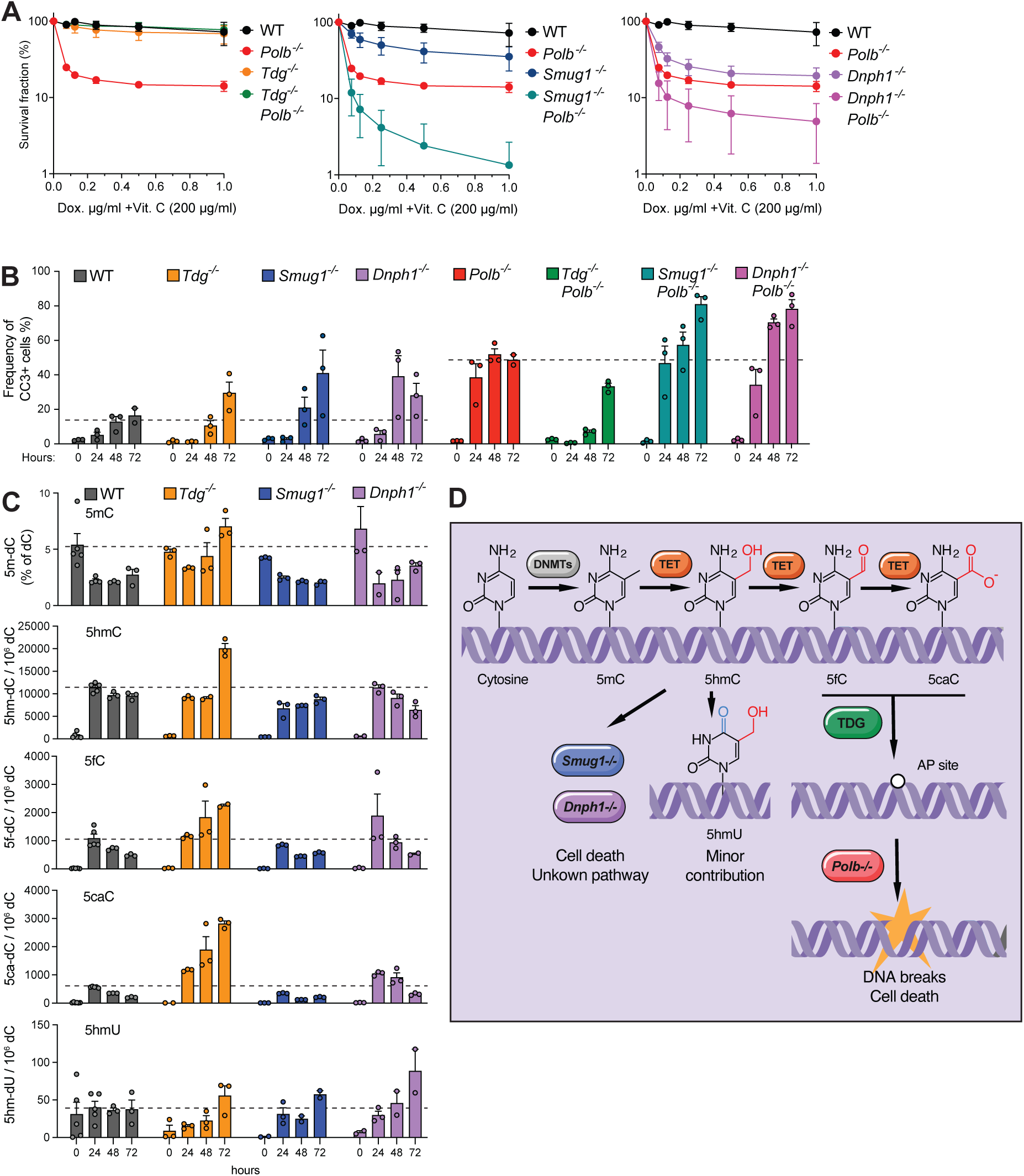
Prolonged TET1 activation drives 5hmU accumulation and sensitivity of *Smug1^−/−^* and *Dnph1^−/−^* cells. **A)** Survival curves of WT-, *Polb^−/−^*-, *Polb^−/−^ Tdg^−/−^*-, *Polb^−/−^ Smug1^−/−^*-, *Dnph1^−/−^*-, *Polb^−/−^ Dnph1^−/−^*-TET1-CD^WT^ clonal cell lines in response to increasing doxycycline concentration and 200µg/mL Vitamin C. Data represents the mean and s.e.m. of n=3 biological repeats (n=3 technical repeats each). **B)** Quantification of apoptosis by cleaved caspase 3 staining and flow cytometry, in response to 1µg/mL doxycycline and 200µg/mL Vitamin C and at different incubation periods. Data represents the mean and s.e.m. of n=3 biological repeats. **C)** Quantification of DNA methylation and oxidation products by LC-MS in WT-, *Tdg^−/−^*-, *Smug1^−/−^*-, and *Dnph1^−/−^*-TET1-CD^wt^ in response to 1µg/mL doxycycline and 200µg/mL Vitamin C at different incubation periods. Data represents the mean and s.e.m. of n=3 biological repeats. **D)** Distinct mechanisms underlie genotoxicity induced by exogenous 5hm-dC and endogenous DNA demethylation. Loss of *Tdg*, but not *Smug1*, rescues DNA damage within 24 h, indicating that oxidative derivatives 5fC/5caC, rather than 5hmU, drive early toxicity in *Polb^−/−^* cells. Over time, 5hmU accumulates in DNA at low levels, while *Smug1^−/−^* and *Dnph1^−/−^* are hypersensitive to oxidative demethylation by an unknown mechanism.

The delayed onset of toxicity raised the possibility that 5hmU could build up at later stages, either through deamination of 5hm-dC in the nucleotide pool or directly on DNA. We performed a time course LC-ID-MS/MS analysis to quantify epigenetic bases over time (**Fig. 7C**). We found a decrease in 5mC and a concomitant increase in oxidised bases 5hmC, 5fC and 5caC, with a larger increase in 5fC and 5caC for the *Tdg^−/−^* cells. 5hmU showed increased levels over time, but its abundance was comparable across all genotypes, with no notable accumulation in *Smug1⁻^/^⁻* and *Dnph1⁻^/^⁻* cells relative to wild type (**Fig. 7C**). We conclude from this that while oxidative demethylation may be a source of 5hmU, particularly at later time points, 5hmU accumulation does not drive cytotoxicity in *Smug1^−/−^* and *Dnph1^−/−^* cells.

Taken together, our results suggest that exogenously supplied 5hm-dC and endogenously generated 5hmC genotoxicity through distinct intermediates (**Fig. 7D**). During the first 24 hours following induction of DNA demethylation, loss of TDG but not SMUG1 fully rescues the phenotype of POLβ-deficient cells, indicating that oxidized cytosine derivatives such as 5fC and 5caC drive genotoxicity during endogenous DNA demethylation. At later time points, however, our observations suggest the presence of an additional, BER-independent and currently unknown mechanism of cytotoxicity associated with oxidative demethylation.

## Discussion

In this study, we directly compared the mechanisms underlying genotoxicity induced by exogenous 5hm-dC and endogenous oxidative DNA demethylation. Although both conditions impose a strong dependency on BER, our data demonstrate that the underlying toxic intermediates are mechanistically distinct. While exogenous 5hm-dC is metabolically converted into 5hm-dU and becomes genotoxic through SMUG1-dependent excision, endogenous TET1-driven demethylation mostly generates toxicity through TDG-mediated excision of 5fC and 5caC. Thus, exogenous and endogenous 5hmC-associated genotoxicity engage BER through distinct upstream intermediates, converging on a shared dependency on repair completion.

For the exogenous genotoxin 5hm-dC, our findings support previous work showing that toxicity arises following its metabolic conversion into 5hm-dU through nucleotide pool deamination, followed by SMUG1-dependent excision (15). Consistent with this model, we observe that toxicity depends on nucleotide metabolism, including DCK-mediated phosphorylation and DCTD-dependent deamination, supporting a pathway in which 5hm-dC is processed prior to DNA incorporation. The requirement for DNPH1 further supports this interpretation, as loss of this sanitisation pathway exacerbates toxicity, likely through accumulation of 5hm-dUMP. Importantly, our data show that loss of *Smug1* fully suppresses DNA damage, cell cycle arrest and apoptosis in POLβ-deficient cells, demonstrating that excision of misincorporated 5hmU is the key step generating cytotoxic BER intermediates in this context. Together, these findings support a model in which genotoxicity arises from misincorporated 5hmU and its subsequent processing, consistent with observations in HR– and FA–deficient systems (14, 15, 18).

In contrast, endogenous oxidative DNA demethylation occurs through a different route. Instead, TET1-induced genotoxicity is dependent on TDG, but not on SMUG1 or DNPH1, indicating that DNA breaks arise early from excision of 5fC and 5caC rather than from deamination-derived 5hmU. While TDG-mediated processing of oxidized cytosines has been shown to generate BER intermediates, including single-strand breaks (11, 12), our data demonstrate that these intermediates can become limiting for cell survival when BER completion is impaired. The lack of involvement of SMUG1 or DNPH1 further argues against a major contribution of nucleotide pool–derived 5hmU, challenging previous work that proposed oxidative demethylation as a source of 5hmU (15). These findings establish that processing of oxidized cytosine derivatives during oxidative DNA demethylation constitutes an intrinsic source of DNA damage that requires efficient downstream repair.

Unexpectedly, however, at later timepoints (48-72 hours) this picture becomes more complex, as SMUG1 and DNPH1 loss confers sensitivity despite the absence of increased 5hm-dU accumulation in these mutants. This effect is largely independent of BER, as the fitness defect persists and is additive in double knockout cells. This suggests the involvement of a currently unknown mechanism operating downstream of, or in parallel to, canonical BER. One possibility is that this reflects the accumulation of non-canonical oxidised species, such as deamination products of 5fC and 5caC (26, 34, 35). Alternatively, epigenetic dysregulation and widespread gene expression alterations may mediate cytotoxicity (36). Together, these observations point to an additional layer of complexity in the cellular response to epigenetic modifications that remains to be defined and warrants further investigation. Oxidative DNA demethylation is particularly prominent in neuronal systems, where extensive TET-driven oxidation coincides with accumulation of TDG-dependent BER intermediates (11, 12). Supporting the importance of BER completion in this context, neurons engage both short-patch and long-patch BER during differentiation, with a substantial contribution from long-patch synthesis at sites of active demethylation, and POLβ-deficient mice exhibit increased neuronal apoptosis and severe developmental defects (37–39), suggesting that failure to resolve demethylation-associated repair intermediates contributes directly to this phenotype.

Beyond the nervous system, TET-dependent oxidation of 5mC also occurs during primordial germ cell (PGC) development, where TET proteins are required for demethylation of imprinted *loci* and meiotic genes through a 5hmC intermediate (40–42). We have recently shown that mice deficient in TLS factors exhibit a severely depleted PGC pool (43), consistent with a requirement for lesion bypass at repair intermediates generated during this demethylation wave. Supporting a broader requirement for replication fork protection in this context, FA-deficient mice also display a PGC depletion phenotype (44), and FANCD2-deficient cells are hypersensitive to exogenous 5hm-dC and its deamination product 5hm-dU, with fork stalling attributed to PARP1 trapping at unresolved BER intermediates (14). We propose that TLS polymerases and FA factors may be required to protect replication forks from collapse at persistent BER intermediates, including AP sites and single-strand breaks arising from TDG-mediated excision of oxidised cytosines, during periods of intense demethylation activity, albeit through distinct mechanisms. The PGC depletion phenotype of TLS- and FA-deficient mice may therefore reflect a common dependency on replication fork integrity during genome-wide demethylation, and our experimental system can provide a tractable platform for identifying essential genes in such contexts. While these findings provide insight into the consequences of oxidative DNA demethylation, they should be interpreted in the context of the experimental system used. Our experimental system relies on expression of the TET1 catalytic domain rather than full-length TET1, which is likely to result in more widespread and less targeted oxidation than occurs under physiological conditions. Nevertheless, the levels of 5hmC induced in our system (10, 000 per 10^6^ cytosines) are comparable to those observed in neurons, where 5hmC reaches its highest known physiological levels in any mammalian cell type (31–33), demonstrating that the modification burden in our system falls within a physiologically relevant range. Although the spatial regulation of TET1 activity in our system may not fully recapitulate endogenous targeting, the overall modification levels are grounded in physiological levels.

In summary, our study demonstrates that exogenous and endogenous 5hmC-associated genotoxicity converge on a shared dependency on BER, yet engage this pathway through distinct upstream intermediates. These findings establish oxidative DNA demethylation as an intrinsic source of DNA damage and highlight the importance of BER capacity in maintaining genome stability during epigenetic reprogramming. In addition, the delayed onset of fitness reduction in a BER-independent manner points to a potential additional layer of genome instability associated with oxidised cytosine derivatives that will require further investigation.

## Acknowledgements

We thank KJ Patel and Julian Sale for sharing published DT40 DNA repair mutants. We thank members of the Garaycoechea lab for critical reading of the manuscript. The research was supported by Dutch Research Council NWO Grant (project OCENW.M20.113).

## Author contributions

J.S. and H.K.W. performed most of the experiments with support from R.H.; E.S. and M.A.T. performed the mass spectrometric analysis of modified nucleosides; figure preparation, H.K.W., J.S., J.I.G.; study concept and design, G.C., J.I.G.; manuscript J.S., H.K.W., J.I.G. with contributions from all authors.

## Materials and Methods

### Cell lines and reagents

32D cells (RRID:CVCL_0118) were cultured in RPMI 1640 (11875093, Gibco) supplemented with 10% fetal bovine serum (FBS; A5256701, Gibco), 1× Penicillin-Streptomycin (PS; 15070063, Gibco), and 5 ng/mL interleukin-3 (IL-3; 213-13, PeproTech). HEK293 (CRL-1573, ATCC) and HEK293T (CRL-3216, ATCC) cells were maintained in DMEM (10564011, Gibco) supplemented with 10% FBS and 1× PS. DT40 DNA repair mutants (gifted by KJ Patel and Julian Sale), were grown in RPMI 1640 supplemented with 7% FBS, 3% chicken serum (16110082, Gibco), and 1× PS. All cell lines were maintained at 37°C in a humidified atmosphere containing 5% CO₂.

### Generation of knockout lines

sgRNAs targeting genes of interest were cloned into pSPCas9(BB)-2A-GFP (PX458; Addgene #48138), lentiCRISPRv2 hygro (Addgene #98291), or pKLV2-U6gRNA5(BbsI)-PGKpuro2ABFP-W (Addgene #67974). Complementary oligonucleotides were annealed in T4 ligase buffer (IVGN2104, Thermo Fisher Scientific), diluted 1:100, and ligated into BbsI-digested (R3539S, NEB), gel-purified vector for 1 hour at room temperature, followed by overnight ligation. Plasmids were transformed into DH5α competent *E. coli* (18265017, Thermo Fisher Scientific) and isolated by miniprep (27104, Qiagen) or maxiprep (12162, Qiagen). All constructs were verified by Sanger sequencing using a U6 promoter primer (Source Bioscience).

To generate *Polb*⁻^/^⁻ cells lacking antibiotic resistance, wildtype 32D cells were transduced with lentiviral particles produced using pSPCas9(BB)-2A-GFP (PX458; Addgene #48138) carrying an sgRNA targeting the *Polb* locus. GFP-expressing clones were isolated by FACS, and loss of POLβ protein was confirmed by western blotting. Knockouts were additionally validated phenotypically by assessing cell survival in the presence of methyl methanesulfonate (MMS; 129925, Sigma Aldrich).

To generate additional wildtype and *Polb*⁻^/^⁻ knockout lines, 32D cells were transduced with lentiCRISPRv2 hygro (Addgene #98291) or pKLV2-U6gRNA5(BbsI)-PGKpuro2ABFP-W (Addgene #67974) transfer plasmids carrying sgRNAs targeting the gene of interest, with virus generated as described below. sgRNA sequences, transfer plasmid backbones, and parental cell lines are detailed in **Supplementary Table 1**. Following selection, pooled populations were validated by western blotting, RT-qPCR, or sensitivity to genotoxic compounds. Where pooled populations could not be validated, individual clones were isolated by limiting dilution and the targeted locus amplified by PCR (primers in **Supplementary Table 2**) and subjected to Sanger sequencing. Indels were identified and analysed by deconvolution of sequencing traces using DECODR (45).

### Lentiviral transduction for the generation of knockout lines

Lentiviral particles were produced in HEK293T cells seeded at 0.7 × 10⁶ cells per well in 6-well plates. After 24 hours, cells were transfected with 0.8 µg psPAX2 (Addgene #12260), 0.2 µg pMD2.G (Addgene #12259), and 1.2 µg transfer plasmid using 6 µL polyethylenimine (PEI; 306185, Sigma Aldrich) in 200 µL serum-free DMEM. After 24 hours, medium was replaced with 32D-specific medium and virus-containing supernatant was harvested 24 hours later, filtered through a 0.45 µm PVDF membrane, and aliquoted for immediate use or stored at −80°C.

For transduction, 0.5 × 10⁶ 32D cells were resuspended in 3 mL virus-containing medium supplemented with polybrene (1:1, 000; TR-1003, Sigma Aldrich) in 6-well plates and spinoculated at 650 × *g* for 90 minutes at 37°C. Transduction was repeated after 24 hours. After a further 48 hours, cells were transferred to selection medium containing either 0.8 mg/mL hygromycin (ant-hg-1, InVivoGen), 1 mg/mL Geneticin (10131035, Gibco), 20 µg/mL blasticidin (A1113903, Gibco), or 0.5 µg/mL puromycin (A1113903, Gibco), as appropriate. Selection was maintained until non-transduced control populations were completely eliminated. Where required, individual clones were subsequently isolated by limiting dilution in 96-well plates; otherwise, pooled populations were expanded for downstream use.

### Generation of inducible TET1 knock in lines

Doxycycline-inducible, TET1 catalytic domain(CD)-expressing targeting constructs were generated based on sequences described previously (28), utilising the pCW57-MCS1-2A-MCS2 backbone (Addgene #71782). EGFP was cloned into the MCS1 position and TET1 CD^WT^ or catalytic-dead TET1 CD^PM^ (H1672Y/D1674A) constructs were cloned into the MCS2 position. The plasmid was further modified to express Turbo-RFP under control of the hPGK promoter. The entirety of this expression cassette was subsequently cloned into pEN396-pCAGGS-Tir1-V5-2A-PuroR TIGRE donor (Addgene #92142) (46), replacing the osTir1-expressing cassette, and puromycin selection was replaced with neomycin. All plasmids were validated by Sanger sequencing.

32D cells stably expressing TET1 CD^WT^ or TET1 CD^PM^ were generated by co-transfection of the modified TIGRE donor vector with pSpCas9(BB)-2A-Puro (PX459) V2.0 (Addgene #62988) (47), for transient expression of SpCas9 and an sgRNA targeting the mouse TIGRE locus. Prior to transfection, 1 × 10⁶ cells were centrifuged at 100 × *g* for 6 minutes, resuspended in 100 µL ice-cold Solution 11 (20 mM HEPES pH 7.5, 5 mM KCl, 10 mM MgCl₂, 90 mM Na₂PO₄), and combined with 5 µg DNA (2.5 µg TIGRE TET1 CD donor plasmid and 2.5 µg pX459). Electroporation was performed using the Lonza Nucleofector 2b system (AAB-1001, Lonza) with programme E-032. Cells were immediately supplemented with 500 µL antibiotic-free medium containing FBS and 10 ng/mL IL-3 (213-13, PeproTech) and transferred to a single well of a 6-well plate in a total volume of 2 mL. After 24 hours, medium was replaced with complete medium supplemented with PS, and after a further 24–48 hours, cells were diluted to 5 × 10⁴ cells/mL in complete medium containing 1 mg/mL Geneticin (10131035, Gibco). Following recovery, individual clones were sorted by FACS into 96-well plates based on Turbo-RFP expression. Clones were screened based on Turbo-RFP expression and successful integration at the TIGRE locus was confirmed by PCR across the 5’ and 3’ homology arms using GoTaq G2 DNA Polymerase (M7841, Promega; annealing temperature 60.7°C); primer sequences are provided in **Supplementary Table 3**. Doxycycline-inducible EGFP expression was confirmed by epifluorescence microscopy and flow cytometry.

### Lentiviral library generation and whole genome CRISPR/Cas9 screening

The gRNA library described in Tzelepis et al. (48), comprising 90, 230 gRNAs targeting 18, 424 genes (approximately 5 gRNAs per gene), was cloned into pKLV2-U6gRNA5(BbsI)-PGKpuro2ABFP-W (Addgene #67974), and the lentiviral library was produced in low-passage HEK293T cells (ATCC). Genome-wide CRISPR/Cas9 screens were performed in wildtype and *Polb*⁻/⁻ 32D cells in biological duplicate; the *Polb*⁻/⁻ suppressor screen additionally utilised two independent clones (C5 and D8). Cell populations were maintained throughout at a minimum 300-fold gRNA coverage.

The MOI was determined empirically by transducing wildtype and *Polb*⁻/⁻ cells with titrated volumes of the viral library and measuring the proportion of BFP-positive cells by flow cytometry 24 hours post-transduction. On Day 0, 75 × 10⁶ cells per condition were transduced at a MOI of 0.3, at which greater than 95% of infected cells are predicted to carry a single integration. BFP positivity was monitored by flow cytometry throughout to confirm adequate library coverage. Successfully transduced cells were selected with 1 µg/mL puromycin (A1113903, Gibco) for 48 hours (Days 2–4), followed by one week of recovery (Days 4–11). Cells were split daily and maintained at 60 × 10⁶ cells per condition in T175 flasks.

5hm-dC treatments were administered during Days 11–13, 16–23, and 26–30. Initial doses of 250 µM and 500 µM (Days 11–13) resulted in unexpectedly high cell viability, attributable to differences in culture scale, and the upper dose was consequently increased to 750 µM with an extended treatment window for subsequent rounds. DNA was sampled prior to treatment initiation and two days after each treatment window, at days 11 (T0), 25 (T1), and 32 (T2). gRNA libraries were sequenced as described in Tzelepis et al. (48) using custom indexing primers (**Supplementary Table 4**), and data were analysed with the model-based analysis of genome-wide CRISPR/Cas9 Knockout (MAGeCK) algorithm (49)

Using the MAGeCK algorithm, we compared sgRNA frequencies between untreated and 5hm-dC-treated conditions. To identify consistently depleted candidates, we applied a multi-parameter prioritization framework across all screening conditions by integrating statistical significance (P value), fold-change (population depletion), MAGeCK score (gene essentiality), and false discovery rate (FDR) Ranking genes by essentiality (negative MAGeCK score) and FDR revealed recurrent hits across all conditions. In addition, we utilized the KEGG (Kyoto Encyclopaedia of Genes and Genomics) classification system to explore the broader biological pathways involved in 5hm-dC processing. For each screen condition, the selection of hits for pathway analysis was conducted based on two different stringency conditions: 1) *P* < 0.001, lfc < −1.5 (**Fig. 1C**) and 2) *P* < 0.01, log fold change (lfc) < −1.5 (**Supplementary Fig. 1B**).

### Immunofluorescence microscopy

No. 1.5 coverslips (631-0150, VWR) were coated with Poly-L-lysine solution (P8920, Sigma Aldrich) for 20 minutes in 24-well plates, then allowed to dry completely at 37°C. 1 × 10⁶ 32D cells were seeded per coverslip and centrifuged at 300 × *g* for 10 minutes. For exogenous 5hm-dC analysis, cells were treated with 5hm-dC-supplemented medium at the concentrations specified in the text for 24 hours. For analysis of TET1 CD overexpression, cells were induced with 1 µg/mL doxycycline and 200 µg/mL L-ascorbic acid 2-phosphate sesquimagnesium salt hydrate (Vitamin C; A8960-5G, Sigma Aldrich) for 12 hours.

For detection of γH2A.X and RPA32 foci, cells were fixed with 4% paraformaldehyde (043368, Alfa Aesar) for 20 minutes and washed three times for 5 minutes with PBS supplemented with 0.5 mM MgCl₂ and 0.5 mM CaCl₂ (PBS-S); coverslips were centrifuged at 300 × *g* for 5 minutes during the first wash. Cells were permeabilised with 0.6% Triton X-100 for 10 minutes, washed once with PBS-S for 5 minutes, and blocked with PBS-S/0.6% Triton X-100 containing 5% BSA (A7638, Sigma Aldrich) for 2 hours. Primary antibodies diluted in blocking buffer were applied overnight at 4°C: anti-phospho-Histone H2A.X (Ser139) (1:1, 000; JBW301, Millipore) and anti-RPA32 (1:100; #2208, Cell Signalling Technology). Coverslips were washed three times for 5 minutes in PBS-S/0.1% Tween-20 and incubated with secondary antibodies diluted in blocking buffer for 1 hour at 37°C protected from light: goat anti-rat Alexa Fluor 488 (1:2, 000; A11006, Thermo Fisher Scientific) and goat anti-mouse Alexa Fluor 594 (1:1, 000; A11032, Thermo Fisher Scientific). Coverslips were washed three times for 5 minutes in PBS-S/0.1% Tween-20, stained with 2 µg/mL DAPI in PBS for 10 minutes, and rinsed once in water before mounting with ProLong Gold Antifade Mountant (P36934, Molecular Probes). Slides were cured for 48 hours prior to imaging. Images were acquired on an LSM 780 confocal microscope (Zeiss). DNA damage foci per nucleus were scored blindly.

For detection of PARylation, cells were fixed in ice-cold 100% methanol (xx) for 15 minutes at −20°C and washed three times in PBS for 5 minutes each. Cells were blocked in PBS/2% BSA/0.3% Triton X-100 for 60 minutes at room temperature. Anti-Poly/Mono-ADP Ribose (D9P7Z) rabbit monoclonal antibody (#89190, Cell Signalling Technology) diluted 1:1, 000 in PBS/1% BSA/0.3% Triton X-100 was applied overnight at 4°C. Coverslips were washed three times in PBS for 5 minutes each and incubated with goat anti-rabbit IgG Alexa Fluor Plus 647 (1:1, 000; A32733TR, Thermo Fisher Scientific) diluted in antibody dilution buffer for 1–2 hours at room temperature protected from light. Coverslips were mounted with ProLong Gold Antifade Reagent with DAPI (P36934, Invtirogen) and images were acquired on a Leica TCS SPE 3 confocal microscope (Leica).

### Protein extractions and immunoblots

Whole-cell extracts were prepared by resuspending 1 × 10⁶ cells in 100 µL RIPA buffer and incubating on ice for 10 minutes. For most experiments (**Supplementary Fig. 1, 6, 7**), RIPA buffer contained 25 mM Tris-HCl pH 7.4, 150 mM NaCl, 0.1% SDS, 0.5% sodium deoxycholate, 1% Triton X-100, and 1× protease inhibitor cocktail (11873580001, Roche), and lysates were clarified by centrifugation at 16, 400 × g for 10 minutes at 4°C. For experiments involving *Tdg⁻/⁻* and control cells (**Supplementary Fig. 11**), RIPA buffer was supplemented with 1 mM EDTA, 1 mM NaF, 3 U benzonase (E1014, Sigma Aldrich), and 10 µL 1 M MgCl₂ (10 mM Tris-HCl pH 7.4, 140 mM NaCl), and lysates were additionally sonicated and clarified by centrifugation at full speed for 30 minutes at 4°C. For detection of SMUG1 and POLB in *Smug1⁻/⁻* and control cells, extracts were instead prepared by resuspending cells directly in 2× Laemmli sample buffer (40 mM Tris-HCl pH 6.8, 3.35% SDS, 16.5% glycerol, 0.005% bromophenol blue, 50 mM DTT) at 1 × 10⁴ cells per µL, followed by addition of benzonase and MgCl₂ (both 1:1, 000) and incubation at room temperature for 1 hour to reduce viscosity. All supernatants were stored at −80°C until analysis. Prior to loading, samples were combined with SDS loading buffer containing DTT and heated at 95°C for 5 minutes (or 70°C for 10 minutes for Laemmli-based extracts).

Proteins were resolved on NuPAGE 4–12% Bis-Tris gels (NP0321BOX, Thermo Fisher Scientific) and transferred to either 0.45 µm nitrocellulose membranes (10600015, GE Healthcare; Supplementary figures 1, 6, 7) or 0.45 µm PVDF membranes (IPVH00010, Millipore; Supplementary figure 11). Membranes were blocked in 5% milk powder in TBS/0.1% Tween-20, except where indicated, and incubated with primary antibodies overnight at 4°C with gentle agitation. The following primary antibodies were used: anti-VINCULIN (1:2, 000, ab129002, Abcam; or 1:1, 000, sc-73614, Santa Cruz, diluted in 5% BSA/TBS-T), anti-POLB (1:250, MA5-12086, Invitrogen; or 1:1, 000, 18003-1-AP, Proteintech), anti-SMUG1 (1:100, sc-514343, Santa Cruz), and anti-TDG (1:400, ab154192, Abcam; or 1:500 for Supplementary figure 11). Membranes were washed three times in TBS/0.1% Tween-20 and incubated with HRP-conjugated secondary antibodies for 1 hour at room temperature: swine anti-rabbit IgG (1:3, 000, P0399, Dako), goat anti-mouse IgG (1:5, 000, P0447, Dako), goat anti-rabbit IgG (1:10, 000, 111-035-003, Jackson), or rabbit anti-mouse IgG (1:10, 000, 315-035-003, Jackson), as appropriate. Signal was detected using either ECL Western Blotting Detection reagent (RPN2106, GE Healthcare) captured on X-ray film, or SuperSignal West Maximum Sensitivity substrate (34096, Thermo Fisher Scientific) imaged on an Amersham IQ800 system (Cytiva).

### Cell viability assays

Cell viability was assessed by CellTiter-Glo 2.0 luminescence assay (G9242, Promega). 32D cells were seeded at 2 × 10⁴ cells per well in clear-bottom white 96-well plates (3610, Costar) in the presence of increasing concentrations of compounds as indicated in the text, prepared by serial 1:2 dilution. Plates were incubated for 72 hours at 37°C, after which CellTiter-Glo 2.0 reagent was diluted 1:1 in culture medium and 50 µL added per well following medium removal. Plates were agitated on an orbital shaker to induce cell lysis and incubated for 10 minutes at room temperature to stabilise the luminescent signal prior to measurement. Luminescence was measured on a plate reader and viability expressed as a percentage of the untreated control after blank subtraction.

Cell viability assays (MTS) were alternatively performed using CellTiter 96 AQueous One Solution Reagent (G3582, Promega). Cells were seeded in 96-well plates at 1 × 10³ cells per well and cultured for 5 days in the presence of increasing concentrations of 5hm-dC (N-1070-100, Jena Bioscience), 5hm-dU (S781754, Sigma Aldrich), gemcitabine (G6423, Sigma Aldrich) or as indicated in the text. Viability was assessed by incubation with CellTiter 96 AQueous One Solution Reagent (G3582, Promega) for 3 hours, and absorbance measured at 490 nm. DT40 IC₅₀ values were determined by fitting dose-response curves with linear regression.

### Cell cycle analysis

Cell cycle analysis was performed using the Click-iT EdU Alexa Fluor 647 Flow Cytometry Assay Kit (C10634, Thermo Fisher Scientific) according to the manufacturer’s instructions. 32D cells were cultured in medium supplemented with exogenous 5hm-dC at the concentrations specified in the text for 72 hours; to allow 5hm-dC wash-out, cells were resuspended in 5hm-dC-free medium for 2 hours prior to EdU labelling. For EdU incorporation, cells were resuspended in medium containing 20 µM EdU and incubated for 2 hours at 37°C. Following labelling, 1 × 10⁶ cells were washed with PBS/1% BSA and resuspended in 100 µL 1× Click-iT fixative for 15 minutes at room temperature. Cells were washed with PBS/1% BSA, resuspended in 100 µL 1× Click-iT saponin-based permeabilisation and wash reagent, and 500 µL of Click-iT reaction cocktail (430 µL 1× reaction buffer, 20 µL CuSO₄, 1.2 µL Alexa Fluor 647 azide, 50 µL 1× buffer additive) was added immediately. Cells were incubated for 30 minutes protected from light, washed with 1× saponin-based permeabilisation and wash reagent, and resuspended in 500 µL of the same reagent containing 0.5 µg/mL DAPI. Samples were immediately acquired on an LSR Fortessa flow cytometer (BD Biosciences) and cell cycle phase distributions were analysed using FlowJo v10.10 (BD Biosciences).

### Apoptosis assays

For detection of apoptosis by Annexin V staining, 32D cells were incubated in medium supplemented with exogenous 5hm-dC at the concentrations indicated in the text for 72 hours, or with 1 µg/mL doxycycline and 200 µg/mL Vitamin C for 24 hours. All subsequent steps were performed on ice unless otherwise stated. Cells were harvested, washed in PBS, and resuspended in Annexin V binding buffer (10 mM HEPES, 140 mM NaCl, 2.5 mM CaCl₂, pH 7.4). A 100 µL aliquot was transferred to a clean FACS tube and incubated with 5 µL Alexa Fluor 647 Annexin V conjugate (A23204, Thermo Fisher Scientific) or Alexa Fluor 488 Annexin V conjugate (A13201, Thermo Fisher Scientific) for 15 minutes at room temperature protected from light. Following incubation, 400 µL Annexin V binding buffer containing 0.5 µg/mL DAPI or 5 µg/mL propidium iodide (PI) was added and the suspension mixed gently. Samples were immediately acquired on an LSR Fortessa flow cytometer (BD Biosciences) and early- and late-stage apoptotic populations were quantified using FlowJo v10.10 (BD Biosciences).

For detection of apoptosis by CC3, 32D cells were resuspended in 100 µL BD Cytofix/Cytoperm solution (554723, BD Biosciences) per well and incubated for 20 minutes at 4°C. Cells were centrifuged at 1, 000 × *g* for 4 minutes and washed twice in 100 µL BD Perm/Wash buffer (554723, BD Biosciences). Cells were centrifuged at 1, 000 × *g* for 4 minutes and resuspended in 50 µL BD Perm/Wash buffer containing primary antibody at 1:500 and incubated for 30 minutes at 4°C in the dark. Cells were centrifuged at 1, 000 × *g* for 4 minutes and washed twice in 100 µL BD Perm/Wash buffer. Cells were centrifuged at 1, 000 × *g* for 4 minutes, resuspended in 50 µL BD Perm/Wash buffer containing goat anti-rabbit IgG Alexa Fluor Plus 647 (1:500; A32733TR, Thermo Fisher Scientific), and incubated for 30 minutes at 4°C protected from light. Cells were centrifuged at 1, 000 × *g* for 4 minutes, washed twice in 100 µL BD Perm/Wash buffer, resuspended in staining buffer, and immediately acquired on a CytFLEX S flow cytometer (Beckman Coulter). Apoptotic populations were quantified using FlowJo v10.10 (BD Biosciences).

### Real-time quantitative PCR and Gene Expression Analysis

Total RNA was extracted from 32D cells using the RNeasy Kit (74104, Qiagen) and first-strand cDNA was synthesised using the QuantiTect Reverse Transcription Kit (205311, Qiagen) according to the manufacturer’s instructions. RT-qPCR for *Dnph1* expression was performed using Brilliant II SYBR Green qPCR Master Mix (600828, Agilent Technologies) on a ViiA 7 thermocycler (Thermo Fisher Scientific) under the following cycling conditions: 50°C for 2 minutes, 95°C for 10 minutes, followed by 40 cycles of 95°C for 15 seconds and 60°C for 1 minute. Mean threshold cycles were determined from three technical replicates using the comparative C_T_ method, and all expression levels were normalised to mouse *Gapdh*. Primer sequences are provided in **Supplementary Table 5**.

### Materials for organic synthesis

Chemicals, unlabeled compounds, enzymes, 3kDa filters for microcentrifugation and solvents were purchased from New England Biolabs, TCI, Sigma-Aldrich, ROTH, Jena Bioscience, MedChemExpress, and Fluorochem and used as received. Acetonitrile and methanol for HPLC-UV and LC-MS analysis were purchased from VWR. The D_3_-5mdC was purchased from LGC Standards. The [^15^N_3_]-5f-dC and [^15^N_3_]-5ca-dC were prepared according to a previously published procedure (29). The [^13^C, ^15^N_2_]-5hm-dU was prepared after treatment with p-formaldehyde according to known procedures in the literature as described below (**Supplementary Fig. 9A**) and the HRMS data confirmed the structure (calc. m/z = 260.0753, found 260.0755) (50). The [^15^N_3_]-5hm-dC was prepared after treatment with NaBH_4_ according to known procedures in the literature (51, 52). DNA samples were spiked with 430 fmol of [^13^C, ^15^N_2_]-5hm-dU, 9 pmol of D_3_-5mdC. In addition, [^15^N_3_]-5ca-dC, [^15^N_3_]-5f-dC and [^15^N_3_]-5hm-dC were added according to the procedure described in the previously published method (29). HPLC and LC-MS columns were purchased from Agilent. Dry glassware and argon atmosphere were used for all water and oxygen sensitive reactions. Demineralized LC-MS/HPLC water (18 MΩ) was used produced by Hydrolab R system. Chromatographic separation and fraction collection were performed using an Agilent 1290 Infinity II LC system (Agilent Technologies, Waldbronn, Germany). Fraction collection was automated using an Agilent 1290 Infinity II Open-Bed Fraction Collector. Instrument control and data acquisition were managed using Agilent OpenLab CDS ChemStation Edition software. The mass spectra and high-resolution mass spectra were recorded on a HESI Thermo Scientific TSQ Quantum Access MAX Triple Quadrupole Mass Spectrometer coupled to an Accela 1250 UHPLC pump and an Accela autosampler and a HESI Q Exactive Focus Orbitrap LC-HRMS System (Thermo Fisher Scientific, Bremen, Germany), respectively. Data were processed using Thermo Scientific Qual Browser in Thermo Xcalibur version 3.063. Reaction progress during the synthesis of the modified nucleosides was monitored by NMR spectroscopy. ^1^H and C NMR spectra were recorded in deuterated (CDCl_3_, CD_3_OD and D_2_O) and nondeuterated solvents on Magritek Spinsolve Carbon 80 MHz spectrometer and processed on SpinSolve or MestReNova software.

### Synthesis of 5-(hydroxymethyl)-2′-deoxyuridine and [^13^C, ^15^N_2_]-5-(hydroxymethyl)-2′-deoxyuridine

The synthesis of 5hm-dUwas achieved based on a slightly modified previously reported procedure (50). Briefly, 2′-deoxyuridine (dU, 1.0 mg, 4 μmol) was placed in a screw capped 4 mL glass vial equipped with a magnetic stirring bar. Paraformaldehyde (2.4 mg, 0.08 mmol) was then added, followed by H_2_O (1 mL) and triethylamine (21.6 μL, 0.144 mmol), corresponding to a molar ratio of 1:20:36 for dU/p-HCHO/Et_3_N. The reaction mixture was then flushed with argon. The vial was heated up at 50 °C and stirred for 17 days. The reaction progress was monitored by TLC and HPLC, revealing a conversion and yield of 55% each. Scaling up to 10 mg dU input under the same conditions and purification of the product on silica allowed product isolation and structure confirmation both by ^1^H and ^13^C NMR spectroscopy, and HRMS analysis (calc. m/z = 257.0779, found 257.0786). Then same reaction conditions were applied to the corresponding isotopically labeled [^13^C, ^15^N_2_]-dU precursor for the synthesis of [^13^C, ^15^N_2_]-5hm-dU and the structure was confirmed by HRMS analysis (calc. m/z = 260.0753, found 260.0755).

### HPLC analysis

The enrichment of 5ca-dC, 5hm-dC, 5hm-dU and 5f-dC after the enzymatic digestion (**Supplementary Fig. 9B**) was carried out on an Agilent 1290 UPLC UV system equipped with a 4.6 × 250mm Agilent Zorbax C18 XBD column (80 Å, 5μm in particle size) thermostated at 20 °C. The chromatographic separation was performed using a gradient elution at a flow rate of 1 mL/min, with 2 mM ammonium formate (pH 8.5) as mobile phase A and acetonitrile as mobile phase B. The gradient started at 100% A and was linearly changed to 99.5% A over 12 min, followed by an isocratic hold at 99.5% A until 62 min. Then it was linearly changed to 80% A over 8 min and to 40% over 5 min. The elution of the unmodified nucleosides was monitored at 260 nm. A typical HPLC trace is depicted in **Supplementary Fig. 9B** and the elution time windows used for the collection and enrichment were: 5.0-6.5 min (5ca-dC), 17-20 min (5hm-dC), 23-25 min (5hm-dU), 35-37 min (5mC) and 66-68 min (5f-dC).

### LC-MS/MS analysis

The LC-MS/MS method parameters used for the analysis of 5mdC, 5hm-dC, 5f-dC, and 5ca-dC and 5hm-dU are reported in **Supplementary Tables 6, 7, 8** and the corresponding calibration curves in **Supplementary Fig. 9C**. Representative LC–MS/MS trace chromatograms of a DNA sample showing the detection of natural and labeled lesions are depicted in **Supplementary Fig. 9D-E**

### NMR and HRMS analysis

The HRMS and ^1^H and ^13^C NMR spectra are reported in **Supplementary Fig. 14.**

### Statistical analyses

The number of independent biological and technical replicates is indicated in the text and figure legends. Unless otherwise stated, data are presented as mean ± s.e.m. and statistical significance was assessed using a two-tailed unpaired t-test. Analysis was performed in GraphPad Prism v10.0.0 (GraphPad Software).

---

**Supplementary Table 1:**
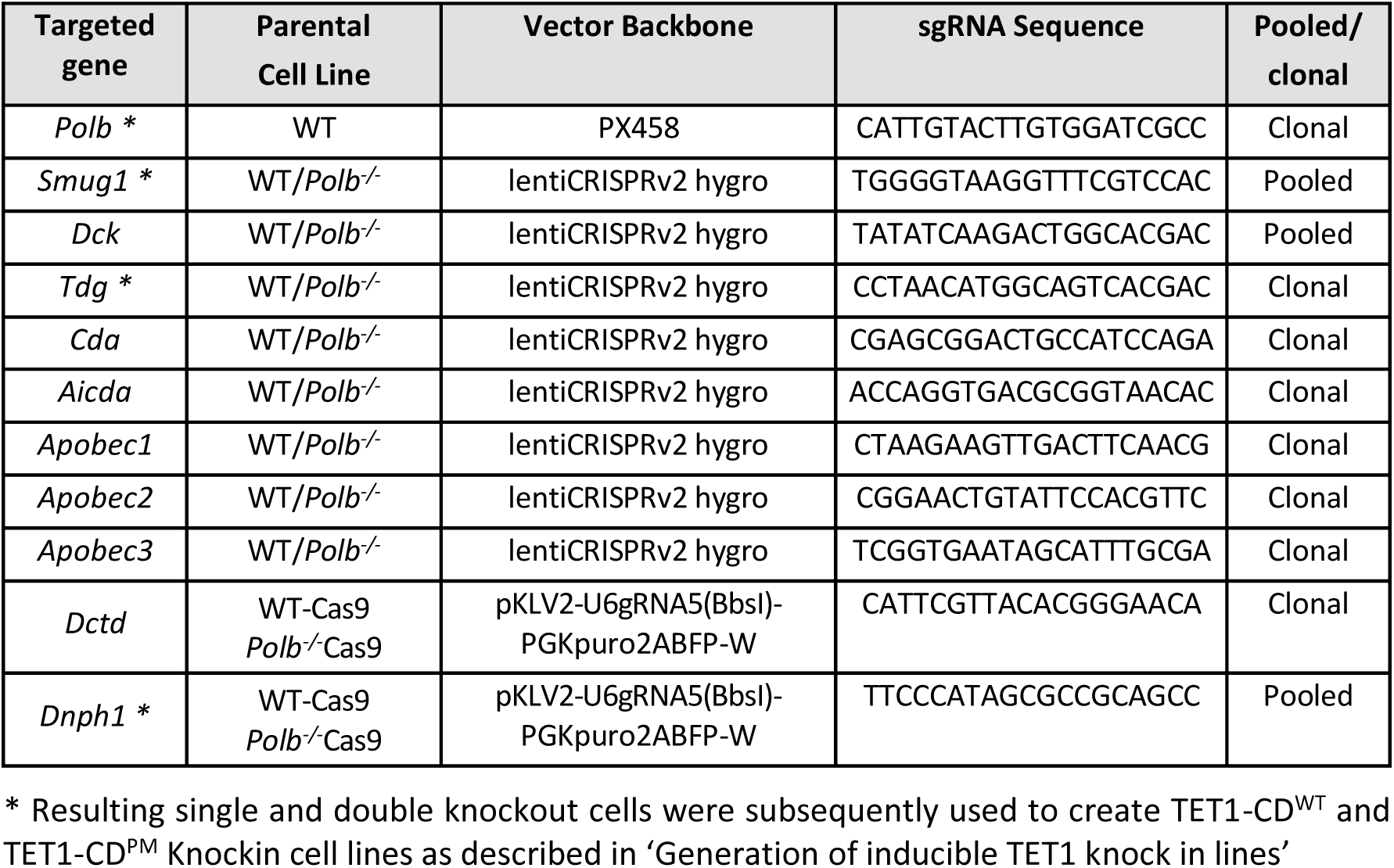
Parental cell lines, plasmid backbones, and sgRNA sequences for the generation of 32D knockout cell lines.

**Supplementary Table 2:**
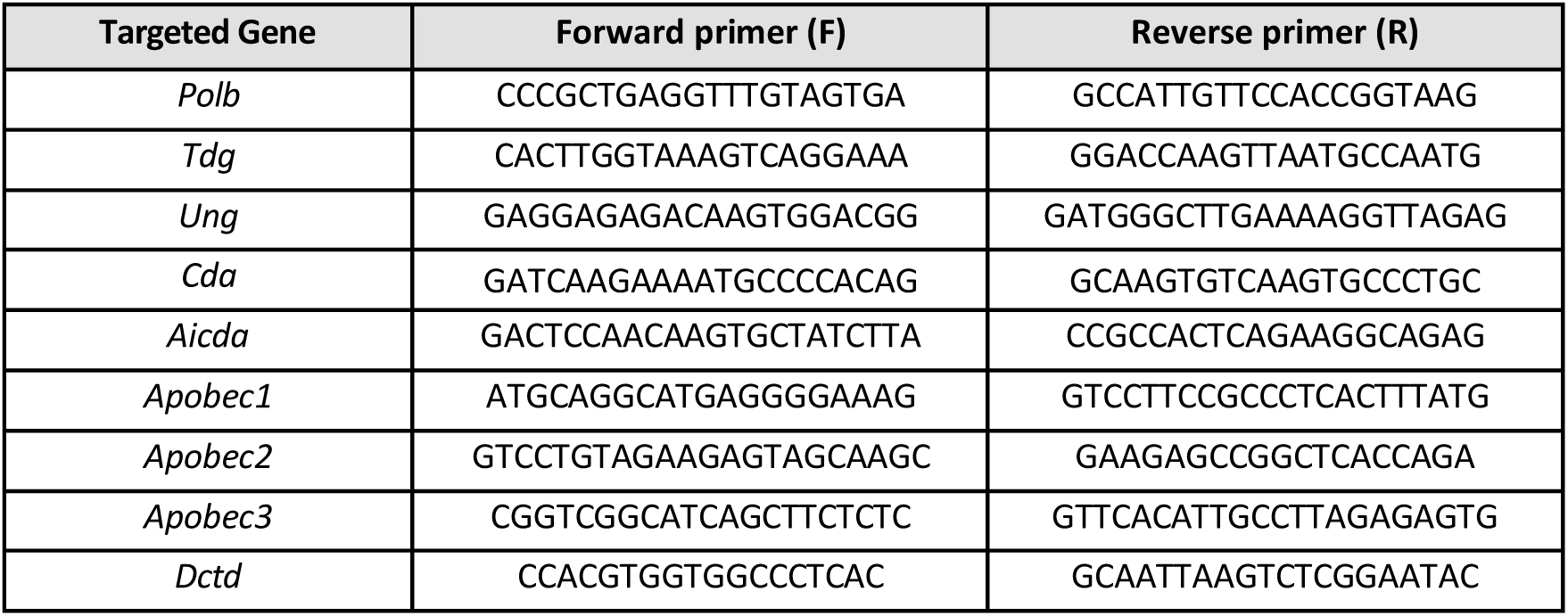
PCR primer for validation of KOs by Sanger sequencing in 32D cells.

**Supplementary Table 3:**
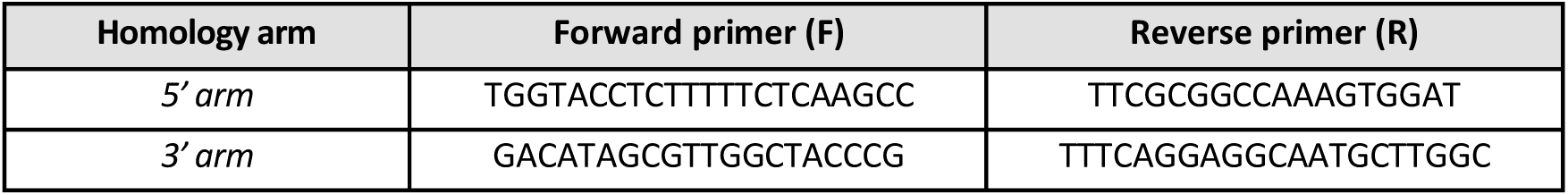
PCR primer sequences for validation of TET1-CD correct integration into the TIGRE locus 32 cells.

**Supplementary Table 4:**
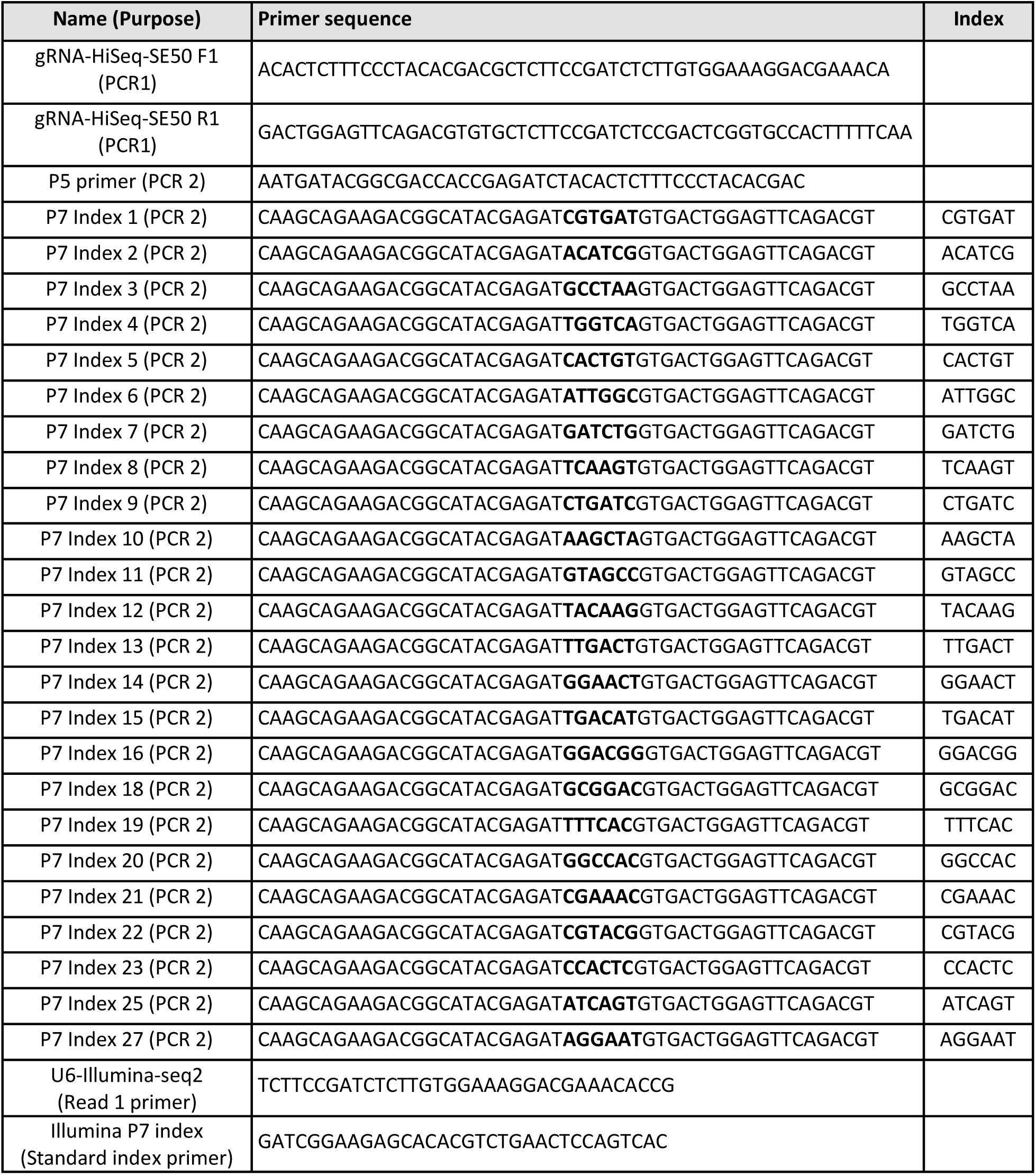
Custom sequencing primers from Tzelepis et al., 2016.

**Supplementary Table 5:**
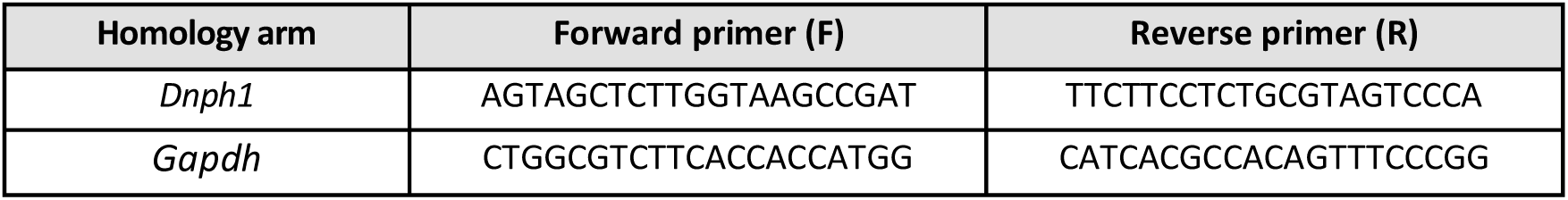
RT-qPCR primers for cultured 32D.

**Supplementary Table 6:** Optimized mass spectrometer parameters for the detection of 5mdC, 5hm-dC, 5hm-dU, 5f-dC and 5ca-dC in SRM mode, including ionization polarity, fragmentation transitions, collision energies, and limits of quantification.

| Analyte | Polarity | MS fragmentation of lesions |  | Optimized parameter | LOQ (fmol) |
| --- | --- | --- | --- | --- | --- |
|  |  | Unlabeled | Isotope-labeled | Collision energy |  |
| 5mdC | + | 242.1 → 126.1 | 245.1 → 129.1 | 15 | 63 |
| 5hm-dC | + | 258.1 → 142 | 261.1 → 145 | 10 | 5.7 |
| 5hm-dU | – | 257.1 → 214 | 260 → 215 | 11 | 5.1 |
| 5f-dC | + | 256.1 → 140 | 259.1 → 143 | 17 | 1.1 |
| 5ca-dC | + | 272.1 → 156.1 | 275.1 → 159.1 | 17 | 0.46 |

**Supplementary Table 7:**
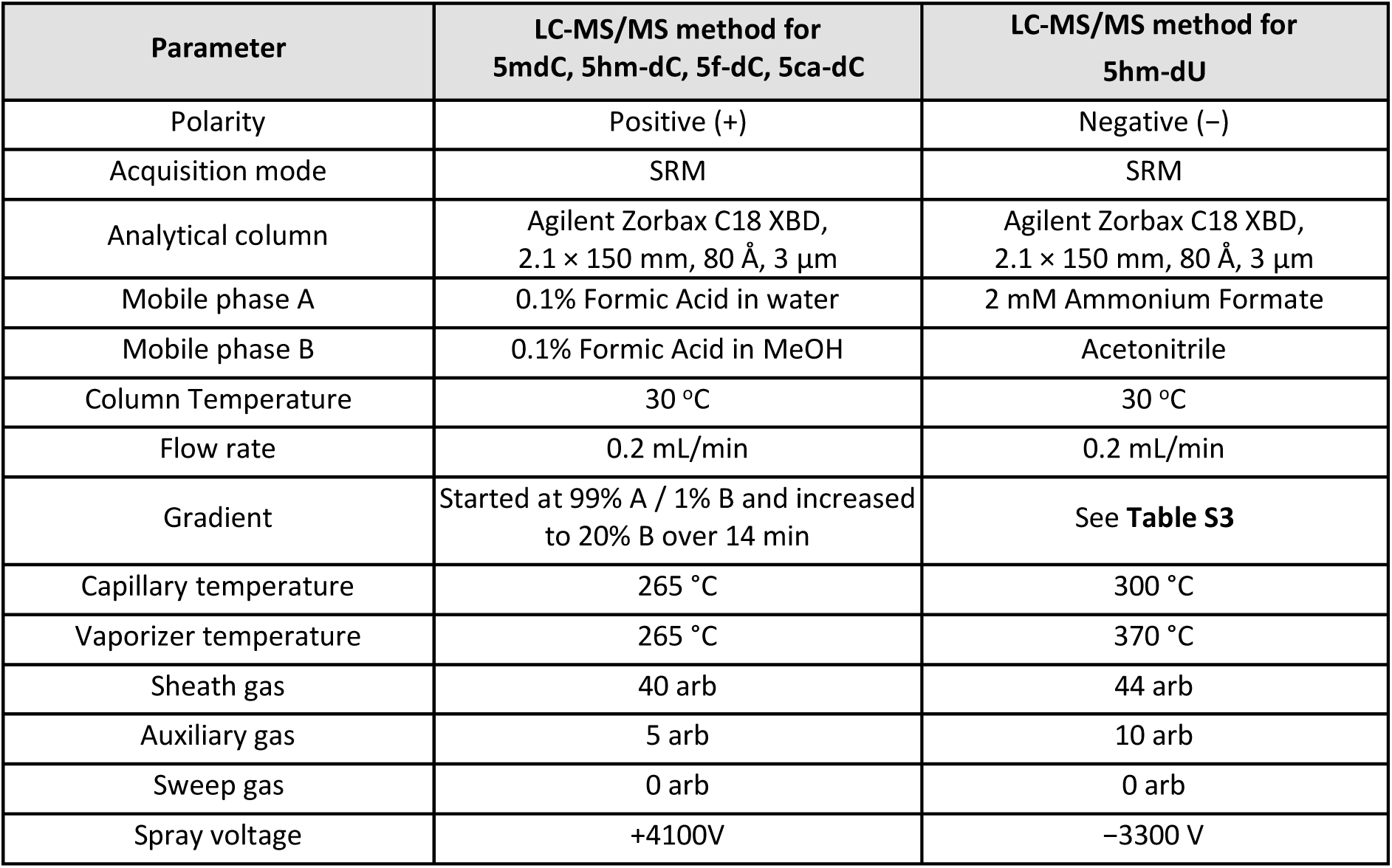
LC-MS/MS method parameters used for the analysis of 5mdC, 5hm-dC, 5f-dC, and 5ca-dC in positive ionization mode and 5hm-dU in negative ionization mode.

**Supplementary Table 8:** Gradient program for the LC-MS/MS method used for 5hm-dU. ^a^ Mobile phase A: ammonium formate 2mM; ^b^ B: acetonitrile.

| Time (min) | A (%)<sup>a</sup> | B (%)<sup>b</sup> |
| --- | --- | --- |
| 0.00 | 97.5 | 2.5 |
| 5.00 | 97.5 | 2.5 |
| 5.10 | 90.0 | 10.0 |
| 8.00 | 90.0 | 10.0 |
| 8.10 | 97.5 | 2.5 |
| 15.00 | 97.5 | 2.5 |

## Notes

### Competing Interest Statement

The authors have declared no competing interest.

## References

1. Kohli, R.M. and Zhang, Y. (2013) TET enzymes, TDG and the dynamics of DNA demethylation. Nature, 502, 472–9.

2. Jones, P.A. and Liang, G. (2009) Rethinking how DNA methylation patterns are maintained. Nat. Rev. Genet., 10, 805–11.

3. Seisenberger, S., Andrews, S., Krueger, F., Arand, J., Walter, J., Santos, F., Popp, C., Thienpont, B., Dean, W. and Reik, W. (2012) The dynamics of genome-wide DNA methylation reprogramming in mouse primordial germ cells. Mol. Cell, 48, 849–62.

4. Wu, H. and Zhang, Y. (2014) Reversing DNA methylation: mechanisms, genomics, and biological functions. Cell, 156, 45–68.

5. Tahiliani, M., Koh, K.P., Shen, Y., Pastor, W.A., Bandukwala, H., Brudno, Y., Agarwal, S., Iyer, L.M., Liu, D.R., Aravind, L., et al. (2009) Conversion of 5-methylcytosine to 5-hydroxymethylcytosine in mammalian DNA by MLL partner TET1. Science, 324, 930–5.

6. Ito, S., Shen, L., Dai, Q., Wu, S.C., Collins, L.B., Swenberg, J.A., He, C. and Zhang, Y. (2011) Tet proteins can convert 5-methylcytosine to 5-formylcytosine and 5-carboxylcytosine. Science, 333, 1300–3.

7. Slyvka, A., Mierzejewska, K. and Bochtler, M. (2017) Nei-like 1 (NEIL1) excises 5-carboxylcytosine directly and stimulates TDG-mediated 5-formyl and 5-carboxylcytosine excision. Sci. Rep., 7, 9001.

8. He, Y.-F., Li, B.-Z., Li, Z., Liu, P., Wang, Y., Tang, Q., Ding, J., Jia, Y., Chen, Z., Li, L., et al. (2011) Tet-mediated formation of 5-carboxylcytosine and its excision by TDG in mammalian DNA. Science, 333, 1303–7.

9. Raiber, E.-A., Beraldi, D., Ficz, G., Burgess, H.E., Branco, M.R., Murat, P., Oxley, D., Booth, M.J., Reik, W. and Balasubramanian, S. (2012) Genome-wide distribution of 5-formylcytosine in embryonic stem cells is associated with transcription and depends on thymine DNA glycosylase. Genome Biol., 13, R69.

10. Krokan, H.E. and Bjørås, M. (2013) Base excision repair. Cold Spring Harb. Perspect. Biol., 5, a012583.

11. Wang, D., Wu, W., Callen, E., Pavani, R., Zolnerowich, N., Kodali, S., Zong, D., Wong, N., Noriega, S., Nathan, W.J., et al. (2022) Active DNA demethylation promotes cell fate specification and the DNA damage response. Science, 378, 983–989.

12. Wu, W., Hill, S.E., Nathan, W.J., Paiano, J., Callen, E., Wang, D., Shinoda, K., van Wietmarschen, N., Colón-Mercado, J.M., Zong, D., et al. (2021) Neuronal enhancers are hotspots for DNA single-strand break repair. Nature, 593, 440–444.

13. Luquette, L.J., Miller, M.B., Zhou, Z., Bohrson, C.L., Zhao, Y., Jin, H., Gulhan, D., Ganz, J., Bizzotto, S., Kirkham, S., et al. (2022) Single-cell genome sequencing of human neurons identifies somatic point mutation and indel enrichment in regulatory elements. Nat. Genet., 54, 1564–1571.

14. Peña-Gómez, M.J., Moreno-Gordillo, P., Narmontė, M., García-Calderón, C.B., Rukšėnaitė, A., Klimašauskas, S. and Rosado, I. V. (2022) FANCD2 maintains replication fork stability during misincorporation of the DNA demethylation products 5-hydroxymethyl-2’-deoxycytidine and 5-hydroxymethyl-2’-deoxyuridine. Cell Death Dis., 13.

15. Fugger, K., Bajrami, I., Silva Dos Santos, M., Young, S.J., Kunzelmann, S., Kelly, G., Hewitt, G., Patel, H., Goldstone, R., Carell, T., et al. (2021) Targeting the nucleotide salvage factor DNPH1 sensitizes BRCA-deficient cells to PARP inhibitors. Science, 372, 156–165.

16. Peña-Gómez, M.J., Suárez-Pizarro, M. and Rosado, I. V (2022) XRCC1 Prevents Replication Fork Instability during Misincorporation of the DNA Demethylation Bases 5-Hydroxymethyl-2’-Deoxycytidine and 5-Hydroxymethyl-2’-Deoxyuridine. Int. J. Mol. Sci., 23.

17. Saxena, S., Nabel, C.S., Seay, T.W., Patel, P.S., Kawale, A.S., Crosby, C.R., Tigro, H., Oh, E., Vander Heiden, M.G., Hata, A.N., et al. (2024) Unprocessed genomic uracil as a source of DNA replication stress in cancer cells. Mol. Cell, 84, 2036–2052.e7.

18. Peña-Gómez, M.J., Rodríguez-Martín, Y., Del Rio Oliva, M., Wijesekara Hanthi, Y., Berrada, S., Freire, R., Masson, J.Y., Reyes, J.C., Costanzo, V. and Rosado, I. V (2025) HMCES corrupts replication fork stability during base excision repair in homologous recombination-deficient cells. Sci. Adv., 11, eads3227.

19. Nabel, C.S., Jia, H., Ye, Y., Shen, L., Goldschmidt, H.L., Stivers, J.T., Zhang, Y. and Kohli, R.M. (2012) AID/APOBEC deaminases disfavor modified cytosines implicated in DNA demethylation. Nat. Chem. Biol., 8, 751–8.

20. Ji, S., Fu, I., Naldiga, S., Shao, H., Basu, A.K., Broyde, S. and Tretyakova, N.Y. (2018) 5-Formylcytosine mediated DNA-protein cross-links block DNA replication and induce mutations in human cells. Nucleic Acids Res., 46, 6455–6469.

21. Takao, N., Kato, H., Mori, R., Morrison, C., Sonada, E., Sun, X., Shimizu, H., Yoshioka, K., Takeda, S. and Yamamoto, K. (1999) Disruption of ATM in p53-null cells causes multiple functional abnormalities in cellular response to ionizing radiation. Oncogene, 18, 7002–9.

22. Vaziri, C., Saxena, S., Jeon, Y., Lee, C., Murata, K., Machida, Y., Wagle, N., Hwang, D.S. and Dutta, A. (2003) A p53-dependent checkpoint pathway prevents rereplication. Mol. Cell, 11, 997–1008.

23. Brown, K.D., Rathi, A., Kamath, R., Beardsley, D.I., Zhan, Q., Mannino, J.L. and Baskaran, R. (2003) The mismatch repair system is required for S-phase checkpoint activation. Nat. Genet., 33, 80–4.

24. Hawn, M.T., Umar, A., Carethers, J.M., Marra, G., Kunkel, T.A., Boland, C.R. and Koi, M. (1995) Evidence for a connection between the mismatch repair system and the G2 cell cycle checkpoint. Cancer Res., 55, 3721–5.

25. Weiner, K.X., Weiner, R.S., Maley, F. and Maley, G.F. (1993) Primary structure of human deoxycytidylate deaminase and overexpression of its functional protein in Escherichia coli. J. Biol. Chem., 268, 12983–9.

26. Masaoka, A., Matsubara, M., Hasegawa, R., Tanaka, T., Kurisu, S., Terato, H., Ohyama, Y., Karino, N., Matsuda, A. and Ide, H. (2003) Mammalian 5-formyluracil-DNA glycosylase. 2. Role of SMUG1 uracil-DNA glycosylase in repair of 5-formyluracil and other oxidized and deaminated base lesions. Biochemistry, 42, 5003–12.

27. Zeng, H., Horie, K., Madisen, L., Pavlova, M.N., Gragerova, G., Rohde, A.D., Schimpf, B.A., Liang, Y., Ojala, E., Kramer, F., et al. (2008) An inducible and reversible mouse genetic rescue system. PLoS Genet., 4, e1000069.

28. Guo, J.U., Su, Y., Zhong, C., Ming, G. and Song, H. (2011) Hydroxylation of 5-methylcytosine by TET1 promotes active DNA demethylation in the adult brain. Cell, 145, 423–34.

29. Stefoudi, E. and Terzidis, M.A. (2026) Palladium-Catalyzed Synthesis of 5-Formyl-2′-Deoxycytidine, 5-Carboxyl-2′-Deoxycytidine, and Nitrogen-15-Labeled Derivatives via Ex Situ-Generated Carbon Monoxide from Chloroform and Application in Isotope Dilution LC–MS/MS Analysis of DNA. ACS Omega, 11, 29015–29025.

30. Blaschke, K., Ebata, K.T., Karimi, M.M., Zepeda-Martínez, J.A., Goyal, P., Mahapatra, S., Tam, A., Laird, D.J., Hirst, M., Rao, A., et al. (2013) Vitamin C induces Tet-dependent DNA demethylation and a blastocyst-like state in ES cells. Nature, 500, 222–6.

31. Kriaucionis, S. and Heintz, N. (2009) The nuclear DNA base 5-hydroxymethylcytosine is present in Purkinje neurons and the brain. Science, 324, 929–30.

32. Szulwach, K.E., Li, X., Li, Y., Song, C.-X., Wu, H., Dai, Q., Irier, H., Upadhyay, A.K., Gearing, M., Levey, A.I., et al. (2011) 5-hmC-mediated epigenetic dynamics during postnatal neurodevelopment and aging. Nat. Neurosci., 14, 1607–16.

33. Wagner, M., Steinbacher, J., Kraus, T.F.J., Michalakis, S., Hackner, B., Pfaffeneder, T., Perera, A., Müller, M., Giese, A., Kretzschmar, H.A., et al. (2015) Age-dependent levels of 5-methyl-, 5-hydroxymethyl-, and 5-formylcytosine in human and mouse brain tissues. Angew. Chem. Int. Ed Engl., 54, 12511–4.

34. Schiesser, S., Pfaffeneder, T., Sadeghian, K., Hackner, B., Steigenberger, B., Schröder, A.S., Steinbacher, J., Kashiwazaki, G., Höfner, G., Wanner, K.T., et al. (2013) Deamination, oxidation, and C-C bond cleavage reactivity of 5-hydroxymethylcytosine, 5-formylcytosine, and 5-carboxycytosine. J. Am. Chem. Soc., 135, 14593–9.

35. Baljinnyam, T., Sowers, M.L., Hsu, C.W., Conrad, J.W., Herring, J.L., Hackfeld, L.C. and Sowers, L.C. (2022) Chemical and enzymatic modifications of 5-methylcytosine at the intersection of DNA damage, repair, and epigenetic reprogramming. PLoS One, 17, e0273509.

36. Bordin, D.L., Lefol, Y., Largartos-Donate, M.J., Montaldo, N.P., Rolseth, V., Tekin, N.B., Lirussi, L., Neurauter, C.G., Liu, Y., Aman, Y., et al. (2025) SMUG1 DNA glycosylase modulates reward behaviour through regulation of olfactory receptor gene expression. Nucleic Acids Res., 53.

37. Kim, J., Kim, K., Mo, J.-S. and Lee, Y. (2020) Atm deficiency in the DNA polymerase β null cerebellum results in cerebellar ataxia and Itpr1 reduction associated with alteration of cytosine methylation. Nucleic Acids Res., 48, 3678–3691.

38. Onishi, K., Uyeda, A., Shida, M., Hirayama, T., Yagi, T., Yamamoto, N. and Sugo, N. (2017) Genome Stability by DNA Polymerase β in Neural Progenitors Contributes to Neuronal Differentiation in Cortical Development. The Journal of Neuroscience, 37, 8444–8458.

39. Sugo, N., Aratani, Y., Nagashima, Y., Kubota, Y. and Koyama, H. (2000) Neonatal lethality with abnormal neurogenesis in mice deficient in DNA polymerase beta. EMBO J., 19, 1397–404.

40. Hackett, J.A., Sengupta, R., Zylicz, J.J., Murakami, K., Lee, C., Down, T.A. and Surani, M.A. (2013) Germline DNA demethylation dynamics and imprint erasure through 5-hydroxymethylcytosine. Science, 339, 448–52.

41. Yamaguchi, S., Hong, K., Liu, R., Shen, L., Inoue, A., Diep, D., Zhang, K. and Zhang, Y. (2012) Tet1 controls meiosis by regulating meiotic gene expression. Nature, 492, 443–7.

42. Gu, T.-P., Guo, F., Yang, H., Wu, H.-P., Xu, G.-F., Liu, W., Xie, Z.-G., Shi, L., He, X., Jin, S., et al. (2011) The role of Tet3 DNA dioxygenase in epigenetic reprogramming by oocytes. Nature, 477, 606–10.

43. Shah, P., Hill, R., Dion, C., Clark, S.J., Abakir, A., Willems, J., Arends, M.J., Garaycoechea, J.I., Leitch, H.G., Reik, W., et al. (2024) Primordial germ cell DNA demethylation and development require DNA translesion synthesis. Nat. Commun., 15, 3734.

44. Hill, R.J. and Crossan, G.P. (2019) DNA cross-link repair safeguards genomic stability during premeiotic germ cell development. Nat. Genet., 51, 1283–1294.

45. Bloh, K., Kanchana, R., Bialk, P., Banas, K., Zhang, Z., Yoo, B.-C. and Kmiec, E.B. (2021) Deconvolution of Complex DNA Repair (DECODR): Establishing a Novel Deconvolution Algorithm for Comprehensive Analysis of CRISPR-Edited Sanger Sequencing Data. CRISPR J., 4, 120–131.

46. Nora, E.P., Goloborodko, A., Valton, A.-L., Gibcus, J.H., Uebersohn, A., Abdennur, N., Dekker, J., Mirny, L.A. and Bruneau, B.G. (2017) Targeted Degradation of CTCF Decouples Local Insulation of Chromosome Domains from Genomic Compartmentalization. Cell, 169, 930–944.e22.

47. Ran, F.A., Hsu, P.D., Wright, J., Agarwala, V., Scott, D.A. and Zhang, F. (2013) Genome engineering using the CRISPR-Cas9 system. Nat. Protoc., 8, 2281–2308.

48. Tzelepis, K., Koike-Yusa, H., De Braekeleer, E., Li, Y., Metzakopian, E., Dovey, O.M., Mupo, A., Grinkevich, V., Li, M., Mazan, M., et al. (2016) A CRISPR Dropout Screen Identifies Genetic Vulnerabilities and Therapeutic Targets in Acute Myeloid Leukemia. Cell Rep., 17, 1193–1205.

49. Li, W., Xu, H., Xiao, T., Cong, L., Love, M.I., Zhang, F., Irizarry, R.A., Liu, J.S., Brown, M. and Liu, X.S. (2014) MAGeCK enables robust identification of essential genes from genome-scale CRISPR/Cas9 knockout screens. Genome Biol., 15, 554.

50. LaFrancois, C.J., Fujimoto, J. and Sowers, L.C. (1998) Synthesis and characterization of isotopically enriched pyrimidine deoxynucleoside oxidation damage products. Chem. Res. Toxicol., 11, 75–83.

51. Münzel, M., Globisch, D., Trindler, C. and Carell, T. (2010) Efficient synthesis of 5-hydroxymethylcytosine containing DNA. Org. Lett., 12, 5671–3.

52. Bienvenu, C., Wagner, J.R. and Cadet, J. (1996) Photosensitized Oxidation of 5-Methyl-2‘-deoxycytidine by 2-Methyl-1, 4-naphthoquinone: Characterization of 5-(Hydroperoxymethyl)-2‘-deoxycytidine and Stable Methyl Group Oxidation Products. J. Am. Chem. Soc., 118, 11406–11411.

