## Supplementayr Figures for "Mechanistically distinct BER processes are essential for the tolerance of exogenous 5hm-dC and enzymatic oxidative demethylation"

1

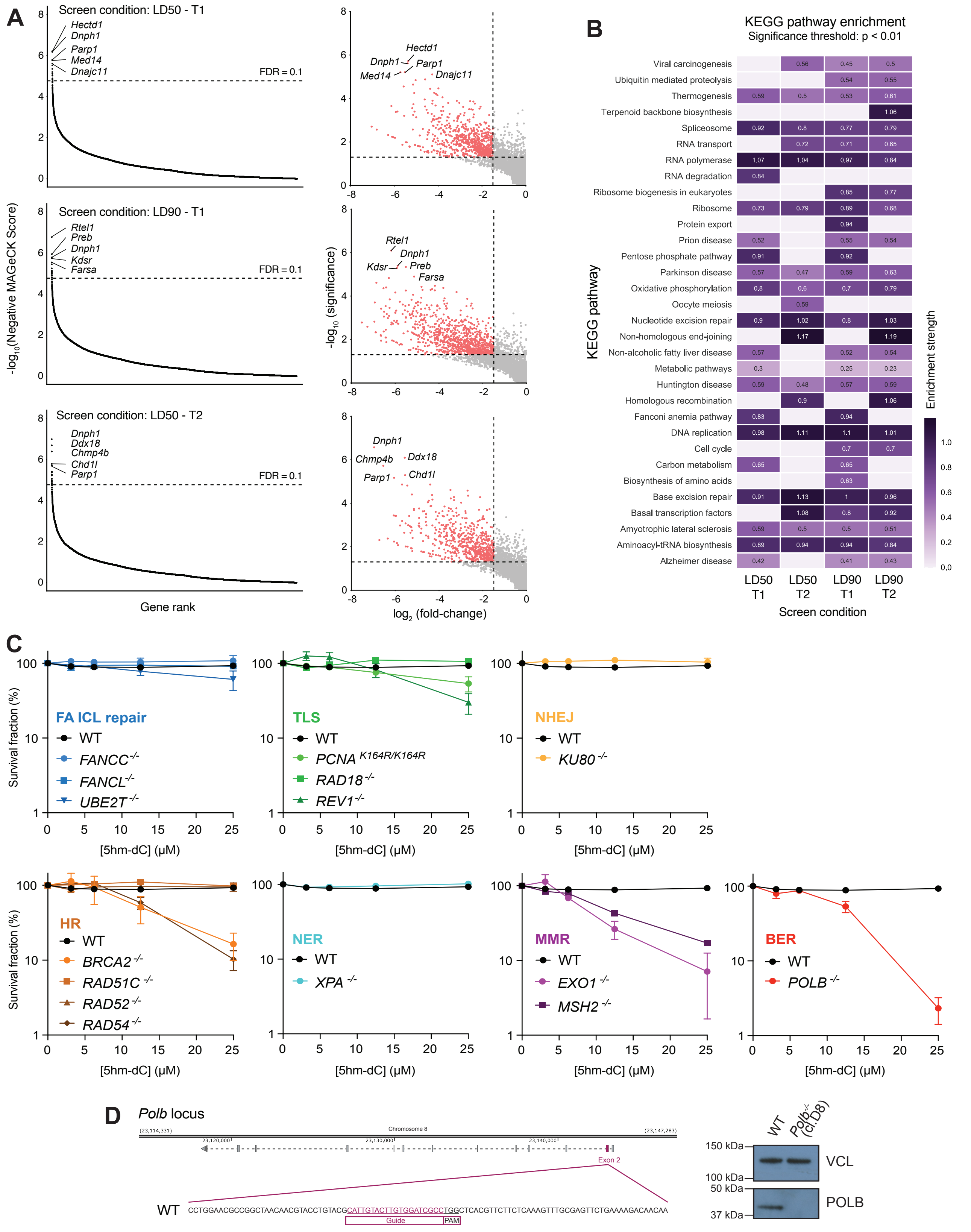


**Supplementary Figure 1. Related to Figure 1. Expanded dataset including additional cell lines, treatment conditions, and full quantification corresponding to the experiments shown in Figure 1. A)** Sensitivity screen results for all 5hm-dC concentrations and timepoints. The left column shows screen results ranked by MAGeCK score, indicating the FDR cutoff (FDR < 0.1). The right column shows volcano plots with the significance scores from MAGeCK analysis. The top five ranked hits according to MAGeCK score for each condition are labelled. **B)** KEGG pathway enrichment analysis of sensitivity screen hits where *P* < 0.01 and log fold change (lfc) < -1.5 for all tested 5hm-dC concentrations and timepoints. Pathway enrichment strength values were determined by calculating Log10(Observed/Expected) for the frequency of proteins appearing in the screen hits versus the expected number of proteins in a given pathway in a random population of factors of the same size **C)** Survival curves of DT40 DNA repair mutants in response to increasing 5hm-dC concentration. Data points represent the mean and s.e.m. of individual biological repeats. (Abbreviation: FA ICL, Fanconi Anaemia Interstrand Crosslink Repair; TLS, Translesion Synthesis; NHEJ, Non-Homologous End Joining; HR, Homologous Recombination; NER, Nucleotide Excision Repair; MMR, Mismatch Repair; BER, Base Excision Repair). **D)** Allele map showing sgRNA targeting *Polb*. Western blot showing loss of *POLB* expression in *Polb*$^{-/-}$ (bottom panel) and vinculin (VCL) loading control (top panel).

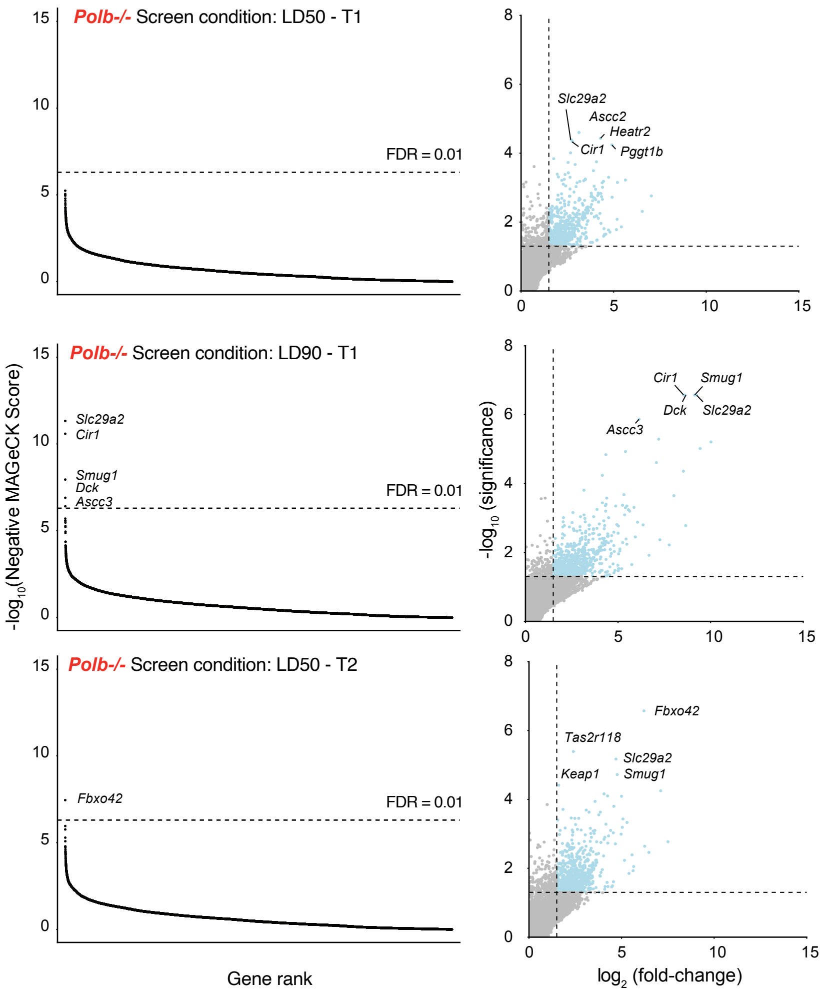


**Supplementary Figure 2. Related to Figure 3. Expanded dataset for the suppressor screen results for all 5hm-dC concentrations and timepoints.** The left column shows screen results ranked by MAGeCK score, indicating the FDR cutoff (FDR < 0.01). The right column shows volcano plots with the significance scores from MAGeCK analysis. The top five ranked hits according to MAGeCK score for each condition are labelled.

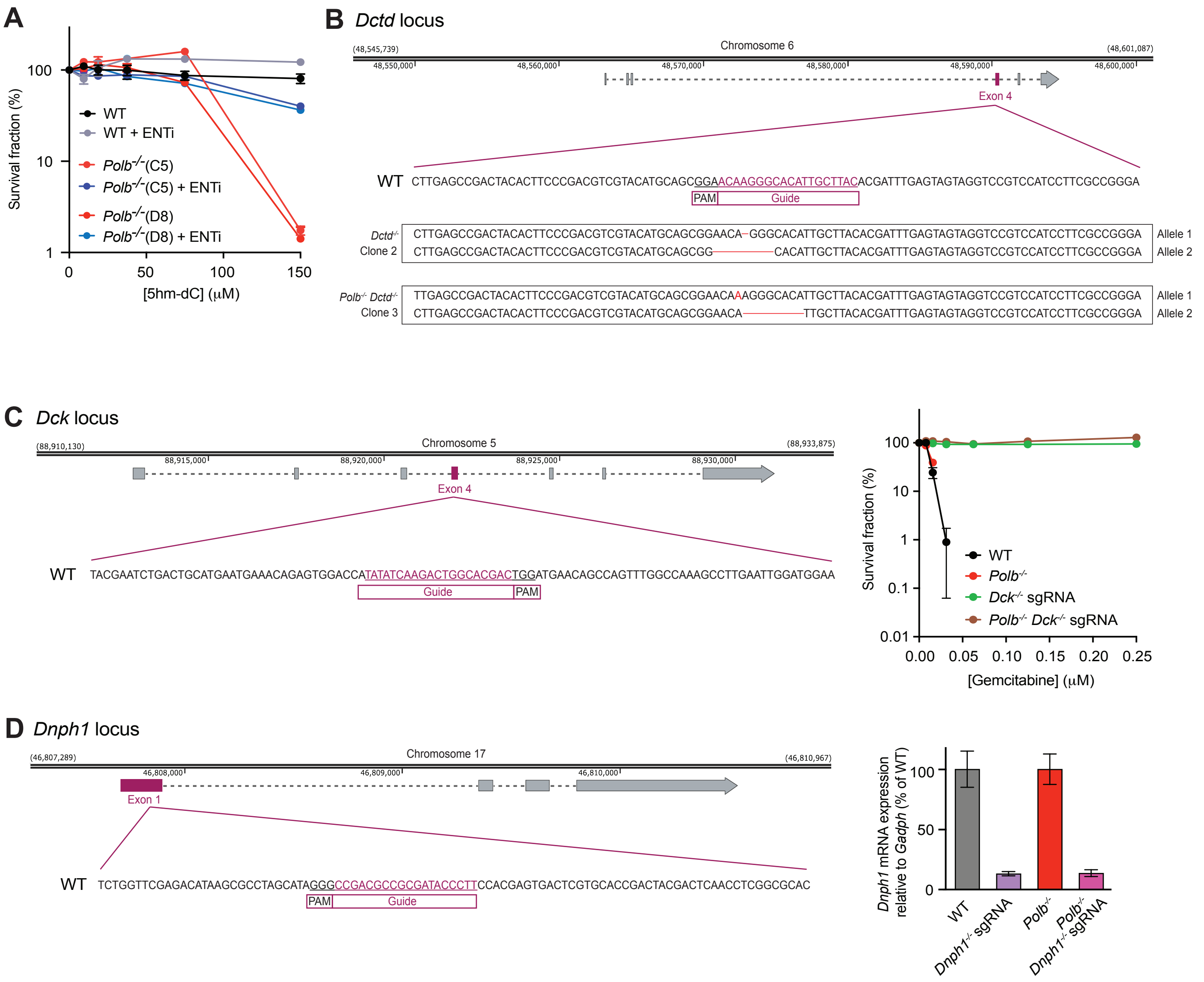


**Supplementary Figure 3. Generation of nucleotide metabolism deficient cells. A)** Survival curves of WT and *Polb*$^{-/-}$ (C5 and D8) clonal cell lines in response to increasing 5hm-dC concentrations in the presence of 18.5μM dipyridamole (ENTi, equilibrative nucleoside transporter inhibitor). Data represents the mean and s.e.m. of n=2 biological repeats. **B)** Allele map showing sgRNA targeting *Dctd*. **C)** Allele map showing sgRN targeting *Dck* and survival curves of *Polb*$^{-/-}$ clonal and *Dck*$^{-/-}$ sgRNA-targeted pooled cell lines in response to increasing Gemcitabine concentration. Data represents the mean and s.e.m. of n=2 biological repeats. **D)** Allele map showing sgRNA targeting *Dnph1* and RT-qCPR validation of loss of *Dnph1* expression in sgRNA-targeted pool cell lines. Fold change normalised to *Gapdh* (relative to parental line). Data represents the mean and s.e.m. of n=3 biological replicates. Corresponding survival curves of WT, *Polb*$^{-/-}$, *Dnph1*$^{-/-}$ sgRNA-targeted pool, and *Polb*$^{-/-}$ *Dnph1*$^{-/-}$ sgRNA-targeted pool cell lines in response to increasing 5hm-dC concentration. Data represents the mean and s.e.m. of n=3 biological repeats (n=3 technical repeats each).

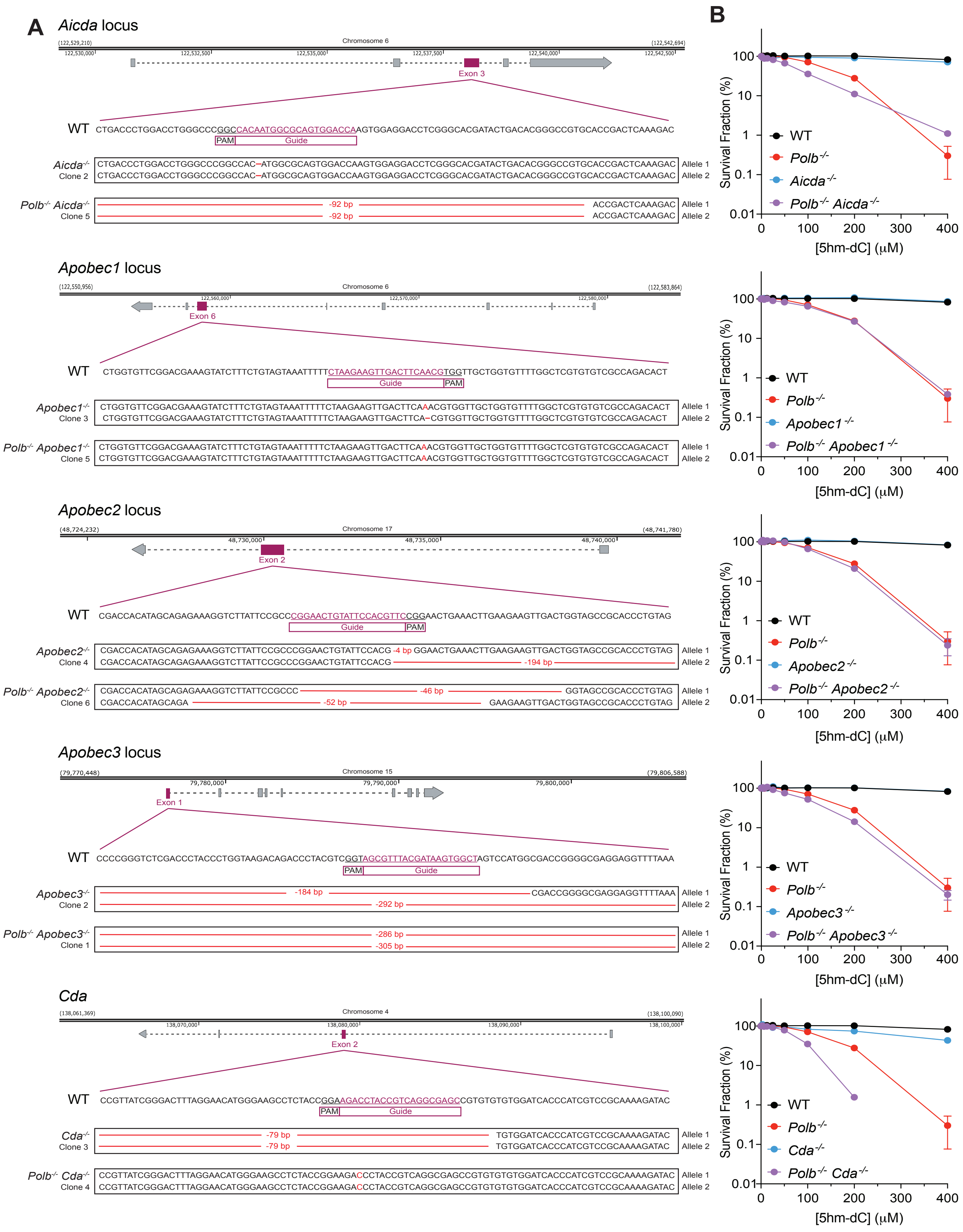


**Supplementary Figure 4. Effects of the loss of different deaminases on WT and *Polb*$^{-/-}$ 32D cell survival in the presence of exogenous 5hm-dC. A)** Allele maps for sgRNAs targeting *Aicda*, *Apobec1*, *Apobec2*, *Apobec3*, and *Cda*. **B)** Survival curves of indicated clonal cell lines in response to increasing 5hm-dC concentration. Data represents the mean and s.e.m. of n=3 biological repeats (n=3 technical repeats each).

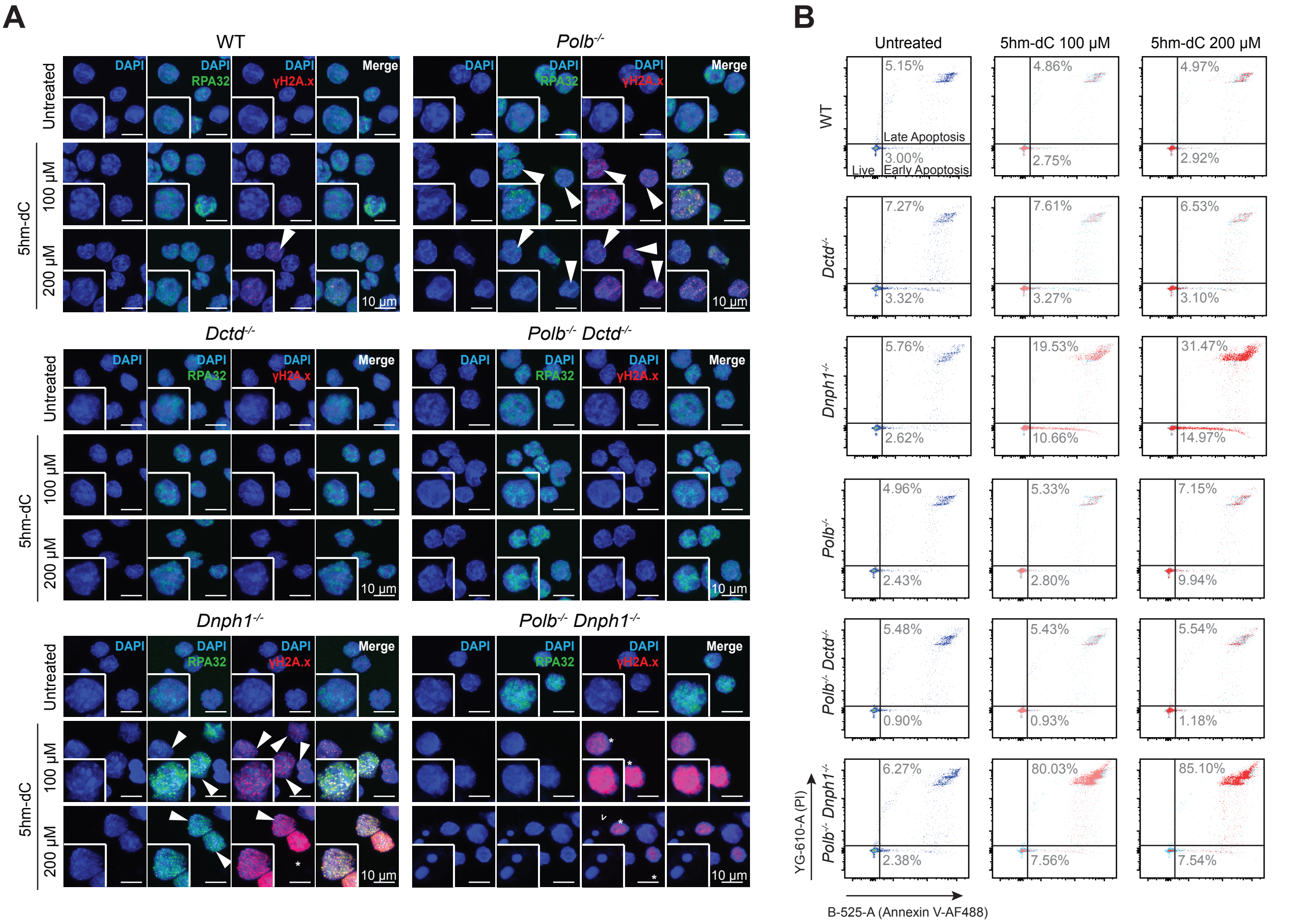


**Supplementary Figure 5. Full set of representative images, quantifications, and knockout validation corresponding to the experiments shown in Figure 3F-G. A)** Full set of representative images of >5 RPA32 or >10 gH2A.X foci in WT, *Polb*$^{-/-}$, *Dctd*$^{-/-}$, *Polb*$^{-/-}$ *Dctd*$^{-/-}$, *Dnph1*$^{-/-}$, *Polb*$^{-/-}$ *Dnph1*$^{-/-}$ clonal cell lines in response to increasing 5hm-dC concentrations. **B)** Full representative flow cytometry plots of early and late apoptotic cell populations in WT, *Polb*$^{-/-}$, *Dctd*$^{-/-}$, *Polb*$^{-/-}$ *Dctd*$^{-/-}$, *Dnph1*$^{-/-}$, *Polb*$^{-/-}$ *Dnph1*$^{-/-}$ clonal cell lines in response to increasing 5hm-dC concentrations. Percentage values represent the mean across experimental replicates.

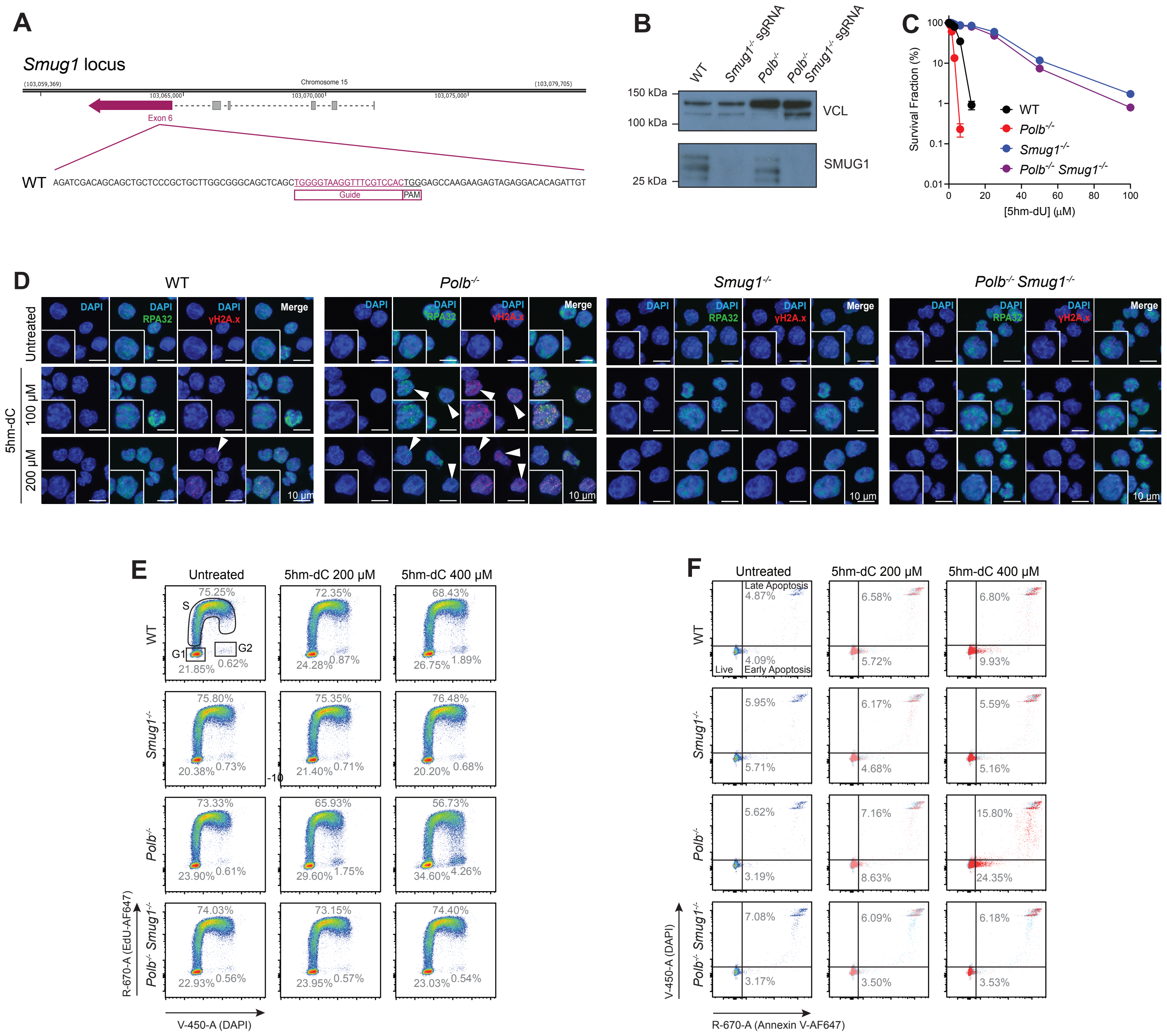

**Supplementary Figure 6. Full set of representative images, quantifications, and knockout validation corresponding to the experiments shown in Figure 4A-D. A)** Allele map showing sgRNA targeting *Smug1* **B)** Western blot of *SMUG1* (bottom panel\and VCL loading control (top panel) in all single and double knockout cell lines. **C)** Survival curves of WT, *Polb*$^{-/-}$, *Smug1*$^{-/-}$ sgRNA-targeted pool, and *Polb*$^{-/-}$ *Smug1*$^{-/-}$ sgRNA-targeted pool cell lines in response to increasing 5hm-dU concentration. Data represents the mean and s.e.m. of n=3 biological repeats (n=3 technical repeats each). **D)** Full set of representative images of >5 RPA32 or >10 gH2A.X foci in WT, *Polb*$^{-/-}$, *Smug1*$^{-/-}$, and *Polb*$^{-/-}$ *Smug1*$^{-/-}$ clonal cell lines in response to increasing 5hm-dC concentrations. **E)** All representative flow cytometry plots of EdU cell cycle analysis in WT, *Polb*$^{-/-}$, *Smug1*$^{-/-}$, and *Polb*$^{-/-}$ *Smug1*$^{-/-}$ clonal cell lines in response to increasing 5hm-dC concentrations. Percentage values represent the mean across experimental replicates and quantification of the percentage of cells in G1, S and G2 phases. **F)** Full representative flow cytometry plots of early and late apoptotic cell populations in WT, *Polb*$^{-/-}$, *Smug1*$^{-/-}$, and *Polb*$^{-/-}$ *Smug1*$^{-/-}$ clonal cell lines in response to increasing 5hm-dC concentrations. Percentage values represent the mean across experimental replicates.

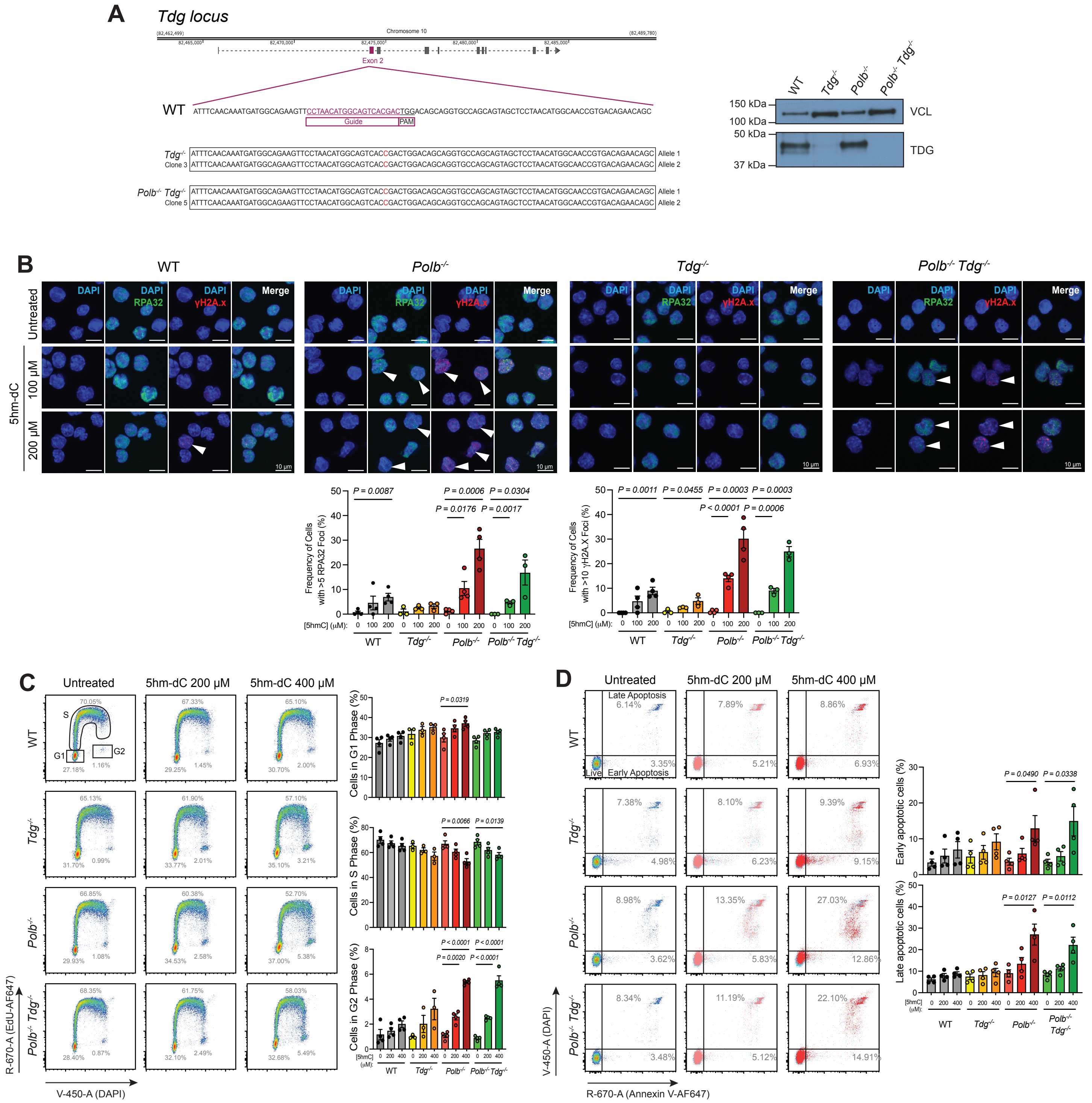

**Supplementary Figure 7. Loss of *Tdg* does not suppress exogenous 5hm-dC induced genotoxicity in *Polb*$^{-/-}$ cells. A)** Allele map showing sgRNA targeting *Tdg* and Western blot of *TDG* (bottom panel) and VCL loading control (top panel) in all single and double knockout cell lines. **B)** Representative images of gH2A.X and RPA32 foci in WT, *Polb*$^{-/-}$, *Tdg*$^{-/-}$, and *Polb*$^{-/-}$ *Tdg*$^{-/-}$ clonal cell lines in response to increasing 5hm-dC concentrations. Arrows indicate cells with >5 RPA32 or >10 gH2A.X foci. Scale bar, 10μM. Corresponding quantification of the percentage of WT, *Polb*$^{-/-}$, *Tdg*$^{-/-}$, and *Polb*$^{-/-}$ *Tdg*$^{-/-}$ clonal cell lines with >5 RPA32 foci (left) and >10 gH2A.X foci (right). Data represents the mean and s.e.m. of n=4 biological replicates for WT and *Polb*$^{-/-}$, and n=3 for *Tdg*$^{-/-}$, and *Polb*$^{-/-}$ *Tdg*$^{-/-}$. *P* values were calculated using an unpaired t-test (only *P* values < 0.05 are displayed). **C)** Representative flow cytometry plots of EdU cell cycle analysis in WT, *Polb*$^{-/-}$, *Tdg*$^{-/-}$, and *Polb*$^{-/-}$ *Tdg*$^{-/-}$ clonal cell lines in response to increasing 5hm-dC concentrations. Percentage values represent the mean across experimental replicates and quantification of the percentage of cells in G1 (top), S (middle) and G2 (bottom) phases in response to indicated 5hm-dC concentrations. Data represents the mean and s.e.m. of n=4 biological replicates. *P* values were calculated using an unpaired *t*-test (only *P* values < 0.05 are displayed) **D)** Representative flow cytometry plots of early and late apoptotic cell populations in WT, *Polb*$^{-/-}$, *Tdg*$^{-/-}$, and *Polb*$^{-/-}$ *Tdg*$^{-/-}$ clonal cell lines in response to increasing 5hm-dC concentrations. Percentage values represent the mean across experimental replicates. Corresponding quantification of the percentage of cells in early (Annexin V-positive, DAPI-negative) and late (Annexin V-positive, DAPI-positive) apoptosis. Data represents the mean and s.e.m. of n=4 biological replicates. *P* values were calculated using an unpaired t-test (only *P* values < 0.05 are displayed).

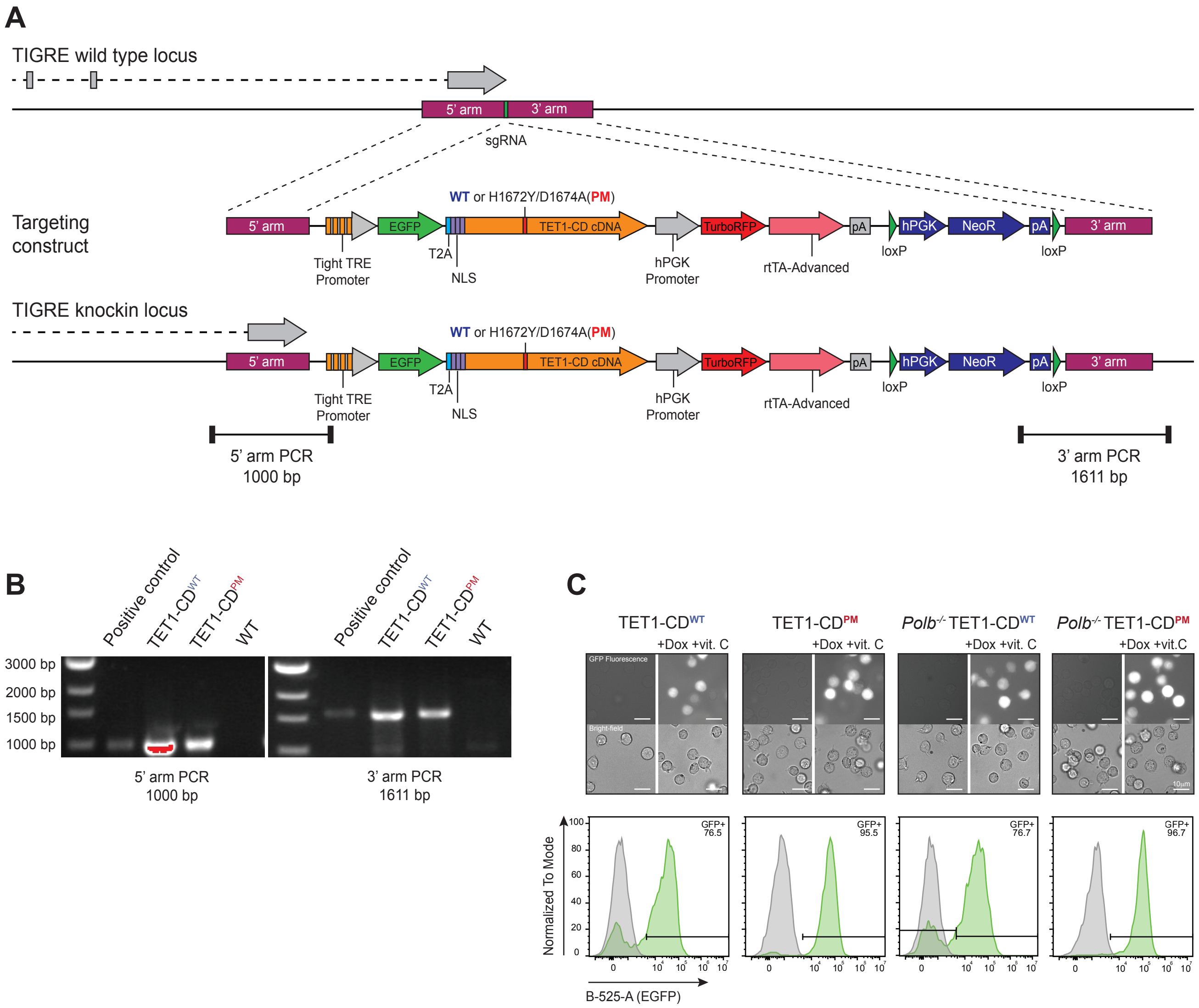


**Supplementary Figure 8. Generation of TET1-CD$^{WT}$ and TET1-CD$^{H1672Y/D1674A(PM)}$ cell lines. A)** Schematic overview of the TET1-CD$^{WT/PM}$ insert. **B)** 5' and 3' Homology Arm PCR of cell lines containing successful integrations. **C)** Representative epifluorescence, brightfield images and flow cytometry plots showing EGFP expression in WT or *Polb*$^{-/-}$ TET1-CD$^{WT/PM}$ cell lines following induction with 1µg/mL doxycycline and 200µg/mL Vitamin C.

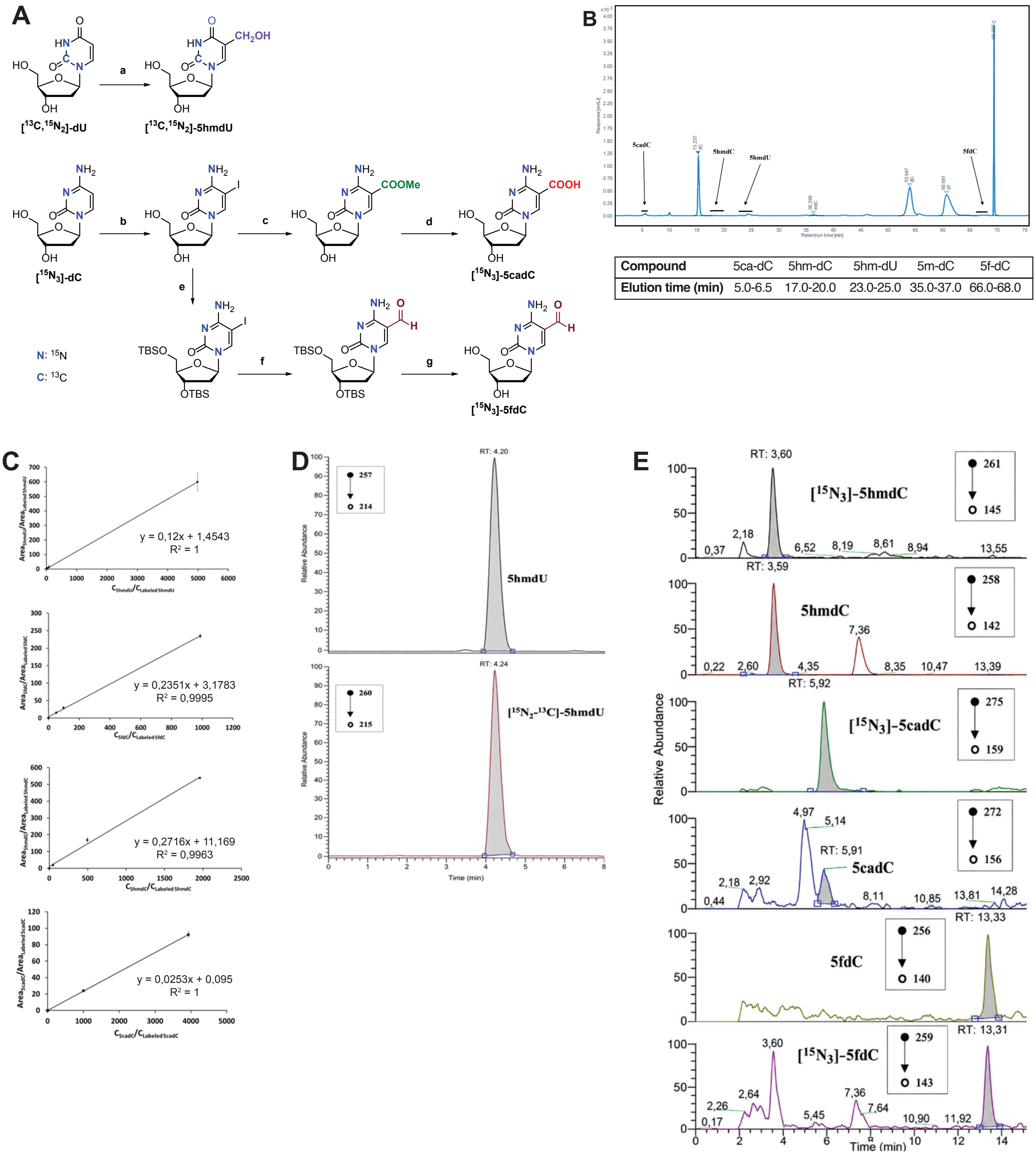


| Compound | 5ca-dC | 5hm-dC | 5hm-dU | 5m-dC | 5f-dC |
|---|---|---|---|---|---|
| Elution time (min) | 5.0-6.5 | 17.0-20.0 | 23.0-25.0 | 35.0-37.0 | 66.0-68.0 |

**Supplementary Figure 9. LC-MS/MS method development and validation for quantification of oxidized DNA bases. A)** Synthesis of [*$^{13}C$, $^{15}N_2$*]-5hmdU, [*$^{15}N_3$*]-5cadC and [*$^{15}N_3$*]-5fdC. (a) *p*-HCHO, Et3N, H2O, 55 °C, 17 days, 55% consumption and conversion; (b) I2, m-CPBA, DMF (dry), 30 min., rt, 64%; (c) vial A: KOH/CHCl3, toluene; vial B: xantphos, Pd2(dba)3, Et3N, methanol, 55 °C, 12 h, 80%; (d) 0.1 M K2CO3, MeOH/H2O 1:1, o.n., 80%; (e) TBS-Cl, imidazole, DMF, 15 h, rt, 65%; (f) vial A: KOH/CHCl3, toluene; vial B: xantphos, Pd2(dba)3, Bu3SnH (6h addition), toluene, 65 °C, 15 h, 50%; (g) TBAF, THF, rt, 50%. **B)** HPLC-UV trace, at 260 nm, of a DNA sample following enzymatic digestion. **C)** Calibration curves employed to quantify 5hmdU, 5hmdC, 5fdC and 5cadC by LC-MS/MS using a stable isotope dilution approach. Plots depict analyte-to-labelled analyte peak area ratios against concentration. Concentrations are expressed in nM. Linear calibration was observed for 5cadC, 5hmdU, 5hmdC and 5fdC; at higher concentrations 5hmdU, 5hmdC and 5fdC exhibited quadratic behaviour (not shown), and the appropriate regression equations were applied where necessary. **D)** LC–MS/MS chromatogram of a representative DNA sample demonstrating detection of 5hmdU and its corresponding [*$^{13}C$, $^{15}N_2$*]-labelled internal standard, acquired in negative ionisation mode. **E)** LC–MS/MS chromatogram of a representative DNA sample demonstrating detection of 5cadC, 5hmdC, 5fdC and their corresponding [*$^{15}N_3$*]-labelled internal standards, acquired in positive ionisation mode.

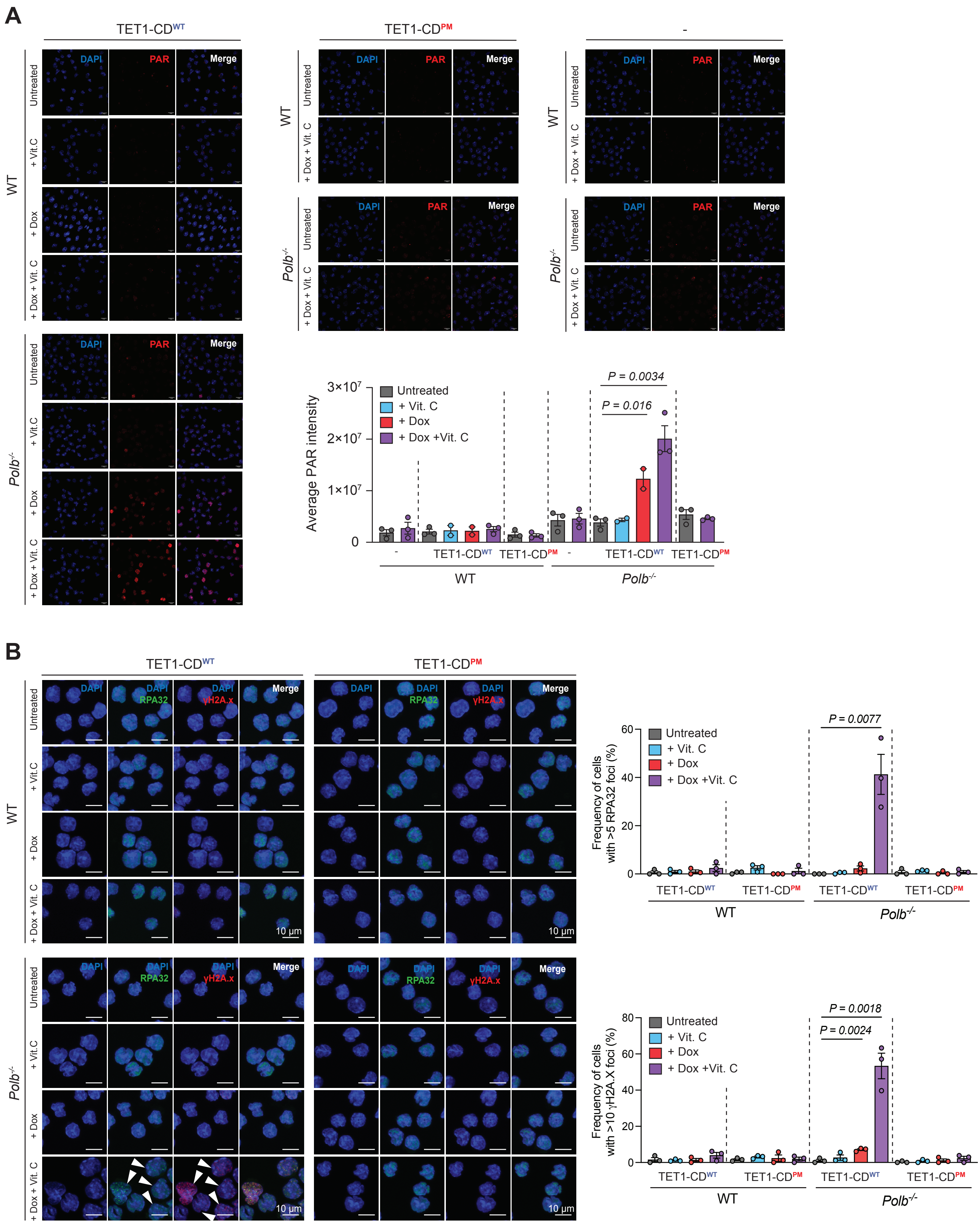


**Supplementary Figure 10. Related to figure 5E-G. Expanded analysis of TET1-induced DNA damage, including additional TET1-CD$^{PM}$ construct. A)** Full set of representative images and quantification of PARylation following TET1 induction, including additional treatment and TET1-CD$^{PM}$ cell lines. **B)** Full set of representative images and quantification of >5 RPA32 or >10 gH2A.X foci following TET1 induction, including additional treatment and TET1-CD$^{PM}$ cell lines.

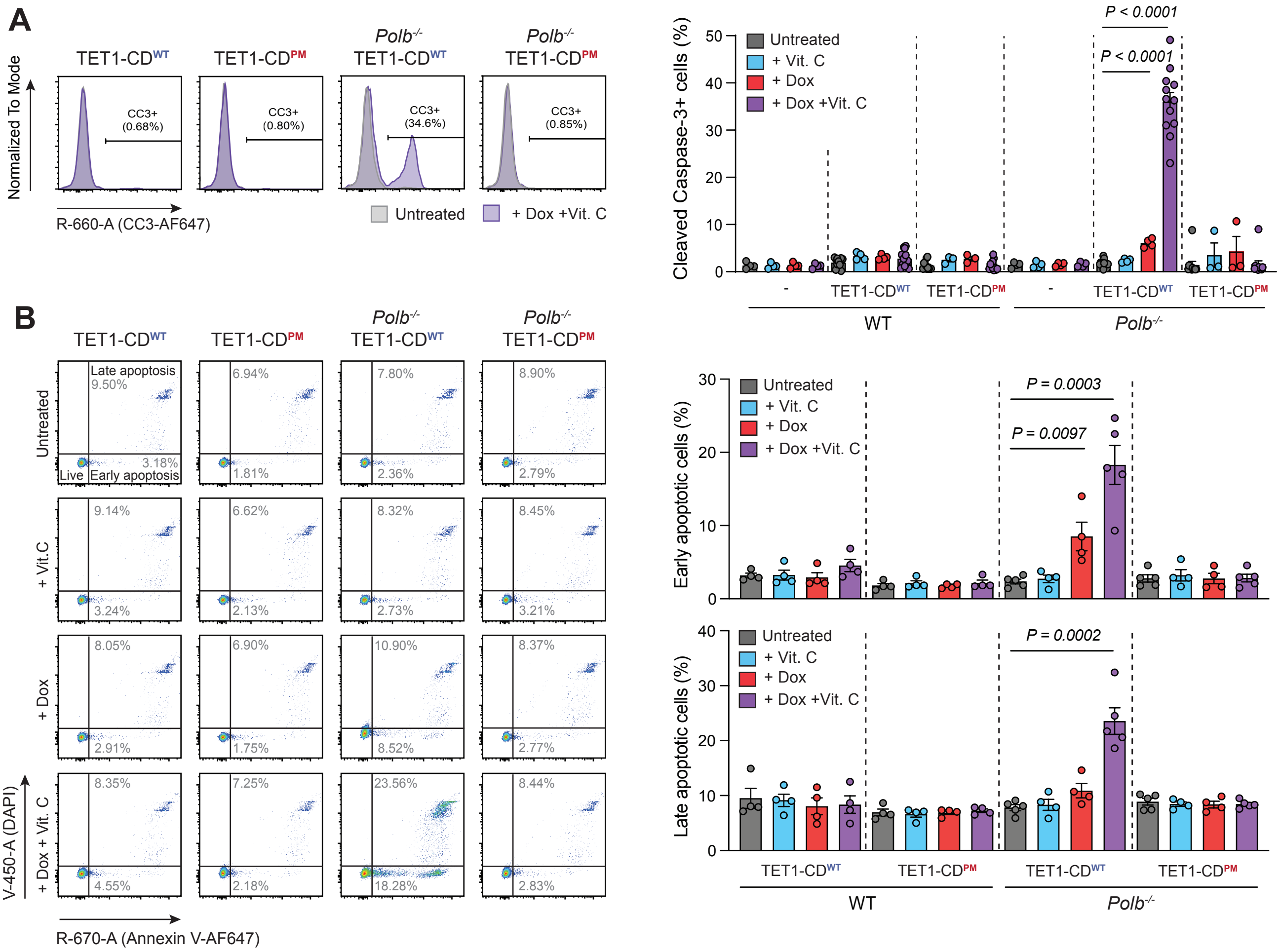


**Supplementary Figure 11. Related to Figure 5H. TET1 mediated oxidative DNA demethylation causes cell death in BER deficient cells. A)** Representative flow cytometry plots and quantifications of apoptotic cell, by cleaved caspase 3, populations following TET1 induction, including additional treatment and TET1-CD$^{PM}$ cell lines. **B)** Representative flow cytometry plots of early and late apoptotic cell populations in WT TET1-CD$^{WT}$, WT TET1-CD$^{PM}$, *Polb*$^{-/-}$ TET1-CD$^{WT}$, and *Polb*$^{-/-}$ TET1-CD$^{PM}$ clonal cell lines following induction with 1μg/mL doxycycline and 200μg/mL Vitamin C. Percentage values represent the mean across experimental replicates. Corresponding quantification of the percentage of cells in early (Annexin V-positive, DAPI-negative) and late (Annexin V-positive, DAPI-positive) apoptosis. Data represents the mean and s.e.m. of n=4 biological replicates. *P* values were calculated using an unpaired t-test (only *P* values < 0.05 are displayed

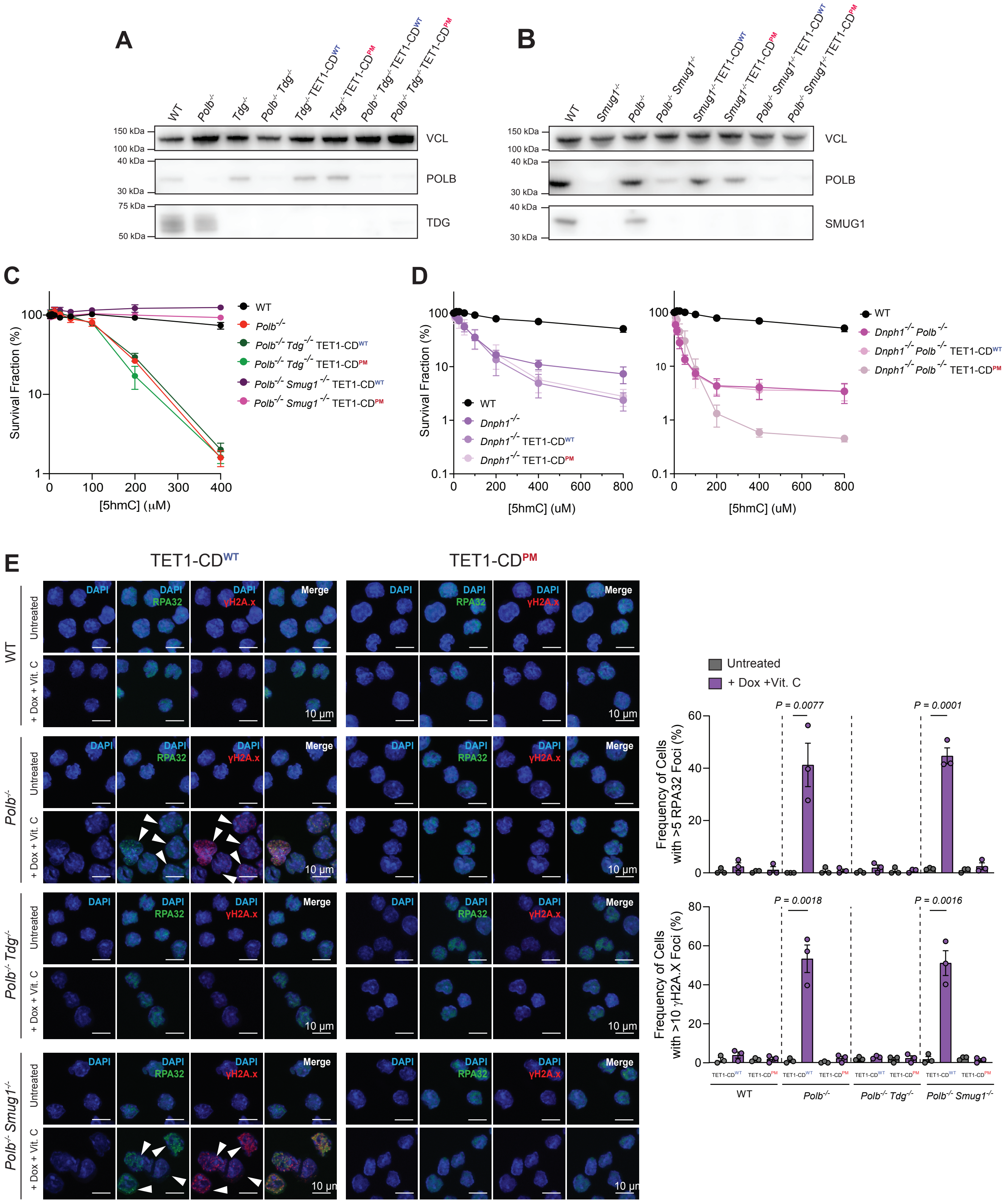


**Supplementary Figure 12. Related to figure 6A-C. Full set of representative images, quantifications, and knockout validation corresponding to the experiments shown in 6A-F. A)** Western blot of *TDG* (bottom panel) *POLB* (middle panel) and VCL loading control (top panel) in all single and double knockout cell lines. **B)** Western blot of *SMUG1* (bottom panel) *POLB* (middle panel) and VCL loading control (top panel) in all single and double knockout cell lines. **C-D)** Survival assay confirmation of sensitivity of cell lines to an increasing 5hm-dC concentration after construct introduction. Data represents the mean and s.e.m. of n=3 biological repeats (n=3 technical repeats each). **E)** Full set of representative images and quantification of >5 RPA32 or >10 gH2A.X foci following TET1 induction, including TET1-CD$^{PM}$ cell lines.

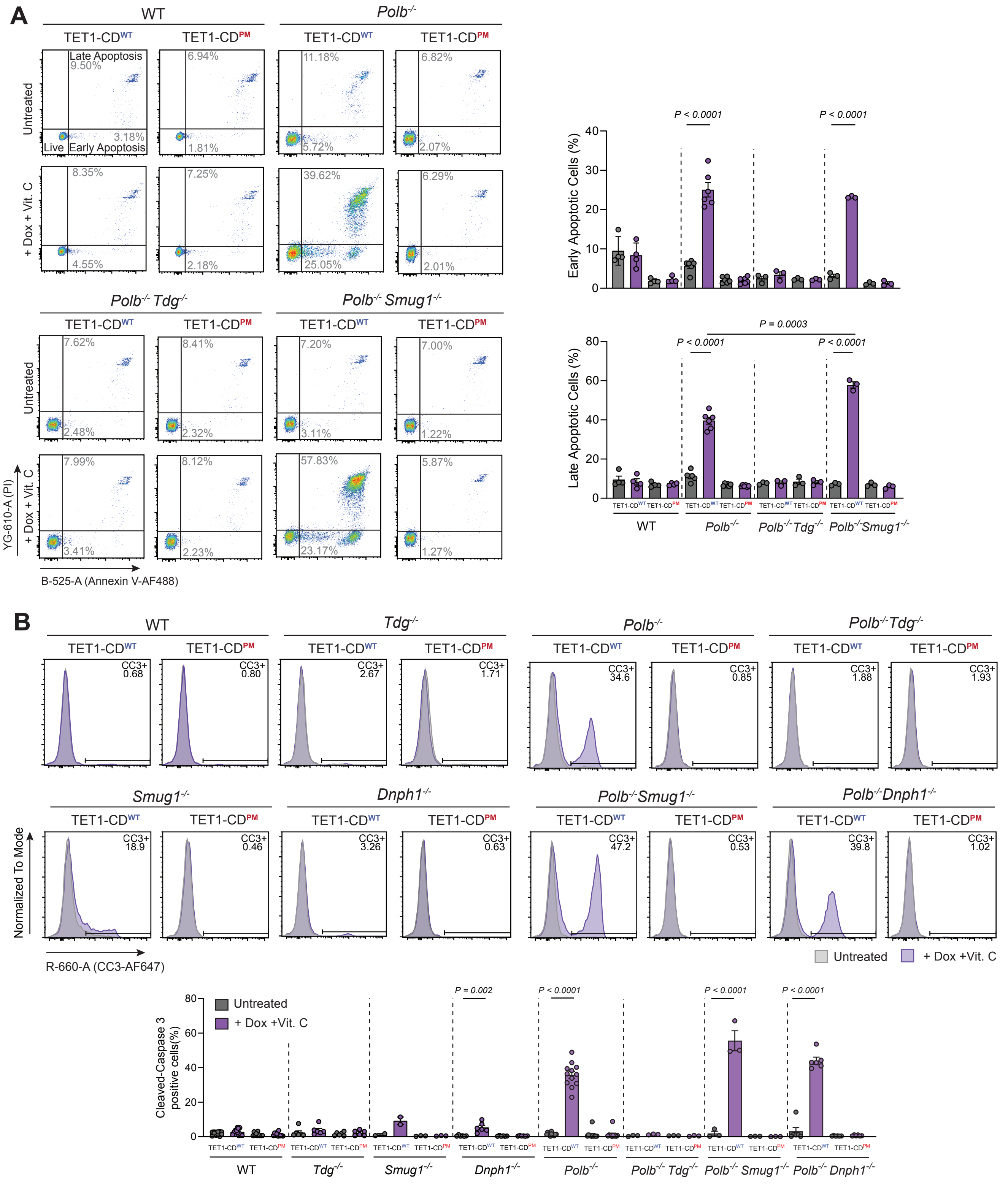


**Supplementary Figure 13. Related to Figure 6D. TET1-mediated cell death is driven by *TDG*-dependent base excision A)** Representative flow cytometry plots of early and late apoptotic cell populations in WT TET1-CD$^{WT}$, WT TET1-CD$^{PM}$, *Polb*$^{-/-}$ TET1-CD$^{WT}$, *Polb*$^{-/-}$ TET1-CD$^{PM}$, *Polb*$^{-/-}$ *Tdg*$^{-/-}$ TET1-CD$^{WT}$, *Polb*$^{-/-}$ *Tdg*$^{-/-}$ TET1-CD$^{PM}$, *Polb*$^{-/-}$ TET1-CD$^{WT}$, and *Polb*$^{-/-}$ *Smug1*$^{-/-}$ TET1-CD$^{PM}$ clonal cell lines following induction with 1µg/mL doxycycline and 200µg/mL Vitamin C. Percentage values represent the mean across experimental replicates. Corresponding quantification of the percentage of cells in early (Annexin V-positive, DAPI-negative) and late (Annexin V-positive, DAPI-positive) apoptosis. Data represents the mean and s.e.m. of n=3biological replicates. *P* values were calculated using an unpaired t-test (only *P* values < 0.05 are displayed). **B)** Representative flow cytometry plots and quantifications of apoptotic cell, by cleaved caspase 3, populations following TET1 induction, including single knockout and TET1-CD$^{PM}$ cell lines.

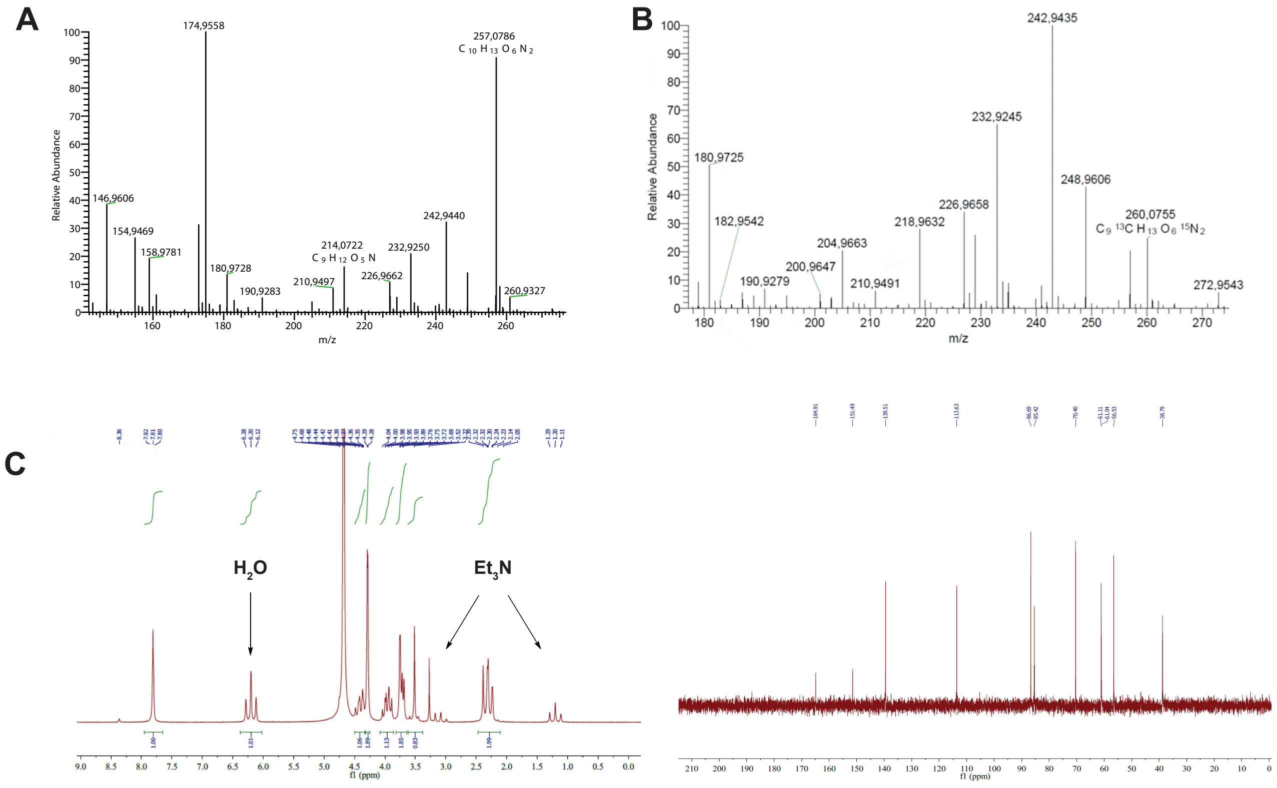


**Supplementary Figure 14. Characterization of synthetic [$^{13}$C, $^{15}N_2$]-5hmdU by high-resolution mass spectrometry and NMR spectroscopy. A)** High-resolution mass spectrum (HRMS) of 5hmdU acquired in negative ion mode, showing the deprotonated molecular ion [M − H]- at *m/z* = 257.0786 and the characteristic daughter ion at *m/z* 214.0722. **B)** HRMS spectrum (ESI, negative ion mode) of [$^{13}$C, $^{15}N_2$]-5hmdU, showing the deprotonated molecular ion [M − H]- at *m/z* = 260.0755. **C)** $^{1}$H and $^{13}$C NMR spectra of 5hmdU.
